# EGFR-to-Src family tyrosine kinase switching in proliferating-DTP TNBC cells creates a hyperphosphorylation-dependent vulnerability to EGFR TKI

**DOI:** 10.1101/2023.07.17.549374

**Authors:** Nazia Chaudhary, Bhagya Shree Choudhary, Anusha Shivashankar, Subhakankha Manna, Khyati Ved, Shagufa Shaikh, Sonal Khanna, Jeetnet Barr, Jagruti Dani, Nandini Verma

## Abstract

Triple-Negative Breast Cancer (TNBC) is the most aggressive type of breast malignancy, with chemotherapy as the only mainstay treatment. TNBC patients have the worst prognoses as a large fraction of them do not achieve complete pathological response post-treatment and develop drug-resistant residual disease. Molecular mechanisms that trigger proliferation in drug-resistant chemo-residual TNBC cells are poorly understood due to the lack of investigations using clinically relevant cellular models. In this study, we have established TNBC subtype-specific cellular models of proliferating drug-tolerant persister (PDTP) cells using different classes of chemotherapeutic agents that recapitulate clinical residual disease with molecular heterogeneity. Analysis of total phospho-tyrosine signals in TNBC PDTPs showed an enhanced phospho-tyrosine content compared to the parental cells (PC). Interestingly, using mass-spectrometry analysis, we identified a dramatic decrease in epidermal growth factor receptor (EGFR) expression in the PDTPs, while the presence of hyper-activated tyrosine phosphorylation of EGFR compared to PC. Further, we show that EGFR has enhanced lysosomal trafficking in PDTPs with a concomitant increase in N-Myc Downstream Regulated-1 expression that co-localizes with EGFR to mediate receptor degradation. More surprisingly, we found that reduced protein levels of EGFR are coupled with a robust increase in Src family kinases, including Lyn and Fyn kinases, that creates a hyper-phosphorylation state of EGFR-Src tyrosine kinases axis in PDTPs and mediates downstream over-activation of STAT3, AKT and MAP kinases. Moreover, paclitaxel-derived PDTPs show increased sensitivity to EGFR TKI Gefitinib and its combination with paclitaxel selectively induced cell death in PDTP-P TNBC cells and 3D spheroids by strongly downregulating phosphorylation of EGFR-Src with concomitant downregulation of Lyn and Fyn tyrosine kinases. Collectively, this study identifies a unique hyper-phosphorylation cellular state of TNBC PDTPs established by switching of EGFR–Src family tyrosine kinases creating a vulnerability to EGFR TKI.

## INTRODUCTION

Triple-negative breast cancer (TNBC) is one of the most aggressive types of epithelial malignancy that is pathologically defined by no or lost expression of estrogen receptor (ER) and progesterone receptor (PR), together with the absence of *ERBB2* amplification (Garrido-Castro, Lin, and Polyak 2019; Penault-Llorca and Viale 2012). TNBC is a highly heterogeneous group of breast malignancies that includes cancer cells of different cellular origins, that is from basal-like, mesenchymal and luminal, and displays a spectrum of histological as well as molecular landscapes with diverse clinical behavior (Hubalek, Czech, and Müller 2017; Yin et al. 2020)(Hubalek, Czech, and Müller 2017; Yin et al. 2020). Despite all these diversities, clinical management of TNBC solely relies on chemotherapeutic agents in different combinations depending on the aggressiveness and metastatic behavior, as TNBC lacks targets for hormone or HER2-directed therapies (Santonja et al., 2018; Yin et al., 2020).

Routine treatment for TNBC patients is cytotoxic neoadjuvant chemotherapy (NACT), followed by postoperative adjuvant chemotherapy (ACT) (Liedtke et al. 2008a, 2008b). The therapy mainly comprises a combination of Taxol base mitotic inhibitors and anthracyclines that intercalate DNA to effectively suppress tumor cell proliferation (Bergin and Loi 2019; Landry, Sumbly, and Vest 2022). Platinum based anti-tumor agents such as cisplatin are combined in metastatic settings. Due to the high mitotic indexes of TNBC tumors, the initial response to NACT is superior in most TNBC compared to the rest of the breast cancer subtypes; however, unfortunately, more than 50% of patients do not achieve pathological complete response (pCR) to NACT and end up with residual disease (RD) which is typically more drug-resistant and metastatic in nature (Dent et al. 2007; Santonja et al. 2018). In different clinical cohorts, the overall and disease-free survival of TNBC patients with RD is very poor and usually less than 5 years (Liedtke et al. 2008a; Zetterlund et al. 2021).

It has been reported that many epithelial tumors including breast, chemo-resistant residual disease, evolve from the primary tumor during the course of therapy due to extensive molecular, metabolic and epigenetic reprogramming (Mikubo et al. 2021; Vallette et al. 2019) (Nedeljković and Damjanović 2019). Several exome sequencing studies on TNBC tumors and patient-derived xenografts (PDXs) suggest that RD rarely exhibits any additional mutational burden (Anand et al., 2021; von Minckwitz et al., 2012). Recently, a large single-cell RNA Sequencing analysis study on chemotherapy-treated patients by Kim et al. (2018) demonstrates that resistance to chemotherapy is mainly adaptive in nature and NACT non-genetically selects residual tumor cells due to transcriptional and molecular reprogramming (Kim et al. 2018). Few genomic studies on Asian TNBC cohorts from Japan and China detect actionable genetic vulnerabilities like alteration in MYC and PTK2 (Kovacevic et al. 2016)(DOI: 10.1200/po.17.00211), while the top altered genes in recurrence in Chinese patients were TP53, PTEN, RB1, PIK3CA and BRCA1 (Nagahashi et al. 2018)(10.1080/07853890.2021.1966086). These studies indicate that Asian TNBC tumors also exhibit genomic makeups not much different from the Western population, as reported in TCGA TNBC data.

Many previous studies have shown cellular models of drug resistance in different types of breast cancer. However, some very recent studies support the conception of residual disease modeled by short-term exposures to cytotoxic drugs that selects a slow-dividing drug-tolerant tumor cell sub-population having the potential to recommence tumor growth after withdrawal of therapy (Abubaker et al. 2013; Blatter and Rottenberg 2015; Kim et al. 2018; Mikubo et al. 2021; Ramirez et al. 2016). Incidentally, most such studies derived from patients and PDX and were focused on genetic or transcriptomic vicissitudes in TNBC, while proteomic and phospho-proteomic reprogramming in response to NACT and during the evolution of proliferating RD is not studied well in TNBC subtypes. Recently, we have reported the development and characterization of proliferating drug-tolerant persister (PDTP) TNBC cells derived from different classes of chemotherapeutics to understand their phenotypic and molecular reprogramming (Chaudhary et al., n.d.)(Chaudhary et al., n.d.)(Chaudhary et al., n.d.)(Chaudhary et al., n.d.) and underlying targetable vulnerabilities. Our study suggests that PDTP state have elevated autophagy dependency and are enriched in mesenchymal phenotypes, and interestingly, these attributes are driven by Glutathione Peroxidase 4 (GPX4) downregulation.

It has been seen that the TNBC PDTPs are more aggressive in terms of proliferation and are resistant to chemotherapy. In order to understand the PDTP state-specific molecular reprogramming in the current study, we have investigated the proteomic changes in TNBC PDTP cells. We identified a dramatic and consistent decrease in epidermal growth factor receptor (EGFR) expression in TNBC PDTPs belonging to different subtypes while a hyper-activation tyrosine phosphorylation of EGFR. Further, we show that EGFR is channelized to the lysosomal pathway in PDTPs due to NDRG1 (N-Myc Downstream Regulated 1) upregulation. Remarkably, we found that EGFR downregulation is coupled with strong upregulation and activation of Src family kinases that creates a hyper-phosphorylation state of the EGFR-Src axle in PDTPs and, thus, over-activates the downstream RTK signals. Lastly, we show that PDTP cells are more susceptible to EGFR TKI Gefitinib. A combination of Paclitaxel and Gefitinib can selectively induce cell death in PDTPs in 2D cultures and 3D spheroids due to a robust collapse of EGFR-Src phosphorylation-dependent cell survival signaling. This study identifies a unique hyper-phosphorylation cellular state in TNBC PDTPs established by switching between EGFR and Src family tyrosine kinase and creating a targetable vulnerability to EGFR TKI that can impede tumor recurrence.

## RESULTS

### Chemotherapy-tolerant proliferating persister TNBC cells undergo EGFR downregulation with a tyrosine hyper-phosphorylation state

To model clinical chemoresistance in terms of residual disease in TNBC, we generated cellular models of PDTPs from representative cell lines from all four subtypes of TNBC [MDA-MB-468, Basal Like -1 (BL1); HCC70, Basal Like-2 (BL-2); HS578T, Mesenchymal Stem-Like (MSL) and MDA-MB-453, Luminal Androgen Receptor positive (LAR) (Chaudhary et al., n.d.)(Chaudhary et al., n.d.)(Chaudhary et al., n.d.)(Chaudhary et al., n.d.). We used cytotoxic doses of three different chemotherapeutic agents, Taxol (paclitaxel), Anthracyclines (doxorubicin), and platinum salt (cisplatin) to select sub-population of residual TNBC cells that sustained viability under therapy and regained the proliferation once the drug was withdrawn, as indicated in the schematic shown in Fig. 1A. The PDTPs were resistant to the last doses of drugs given for selection.

**Figure 1.**
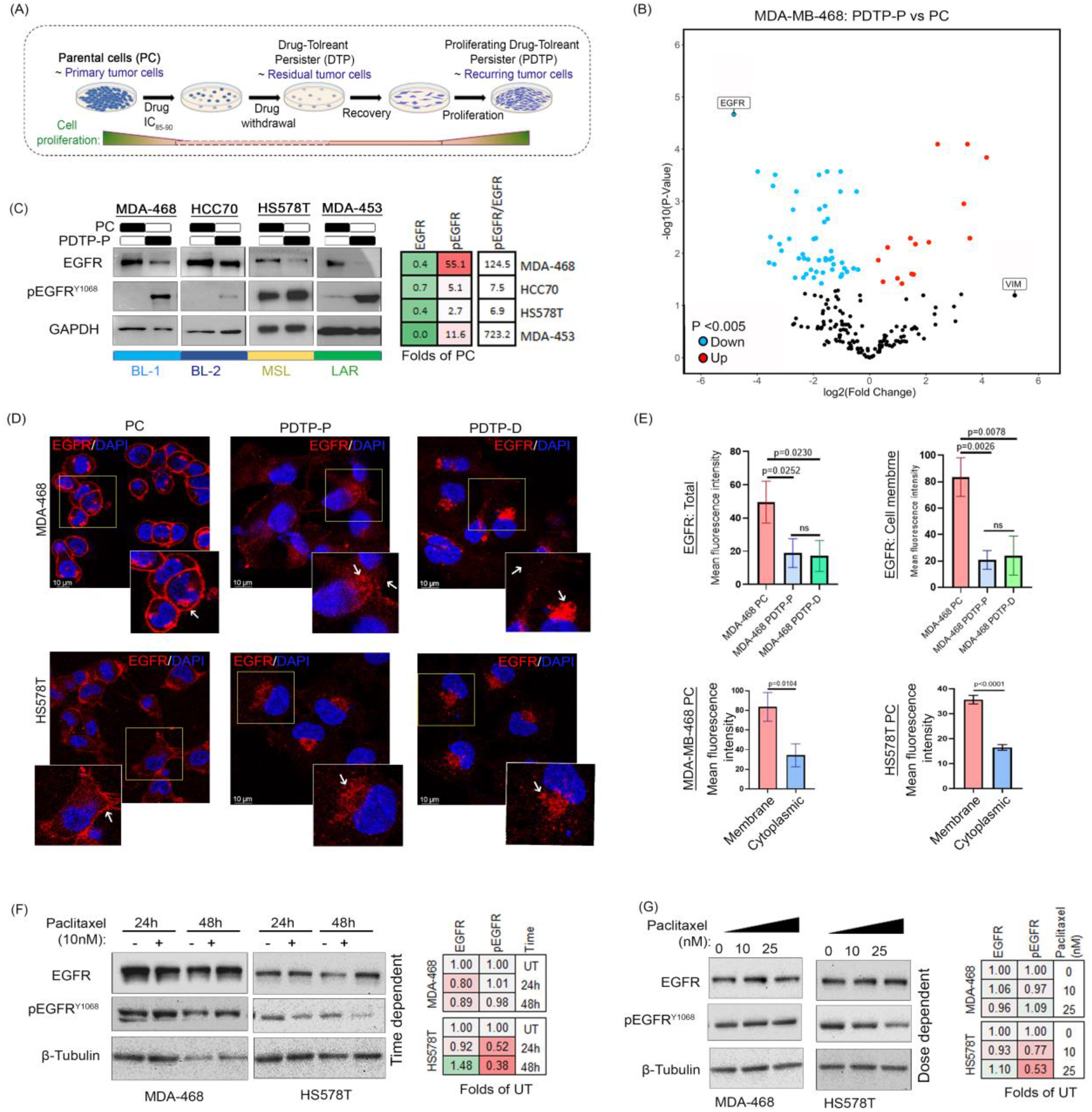
EGFR downregulation and increased tyrosine phosphorylation are associated with PDTP state in all TNBC subtypes. **(A)** Schematic of Proliferating drug-tolerant persister (PDTP) model. Different subtypes of Triple-negative Breast cancer (TNBC) parental cell (PC) lines MDAMB4-68, HCC70, HS478T, and MDA-MB-453 were treated short-term with chemotherapy cytotoxic doses (IC_80-95_) in vitro. After a few days, quiescent drug-tolerant cancer cells were observed. Over time, these quiescent cancer cells resumed growth, establishing “recurrent” colonies. The PDTPs were resistant to the last doses of drugs given for selection. **(B)** The volcano plot below depicts the protein expression levels of 207 proteins as observed by mass spectrometry. The p-values are FDR-adjusted prior to further analysis and the level of significance is set at 0.05. Following these 15 significantly upregulated genes have been plotted in red and 52 significantly downregulated genes have been plotted in blue. EGFR is found to be the most down-regulated. **(C)** Levels of EGFR and p-EGFR (Y^1068^) were determined in different subtypes of parental PC of TNBC and their Paclitaxel-PDTP cells. GAPDH served as their loading control. The expression levels of the indicated proteins were assessed by WB. Band intensities were quantified by ImageJ software and presented as fold of PC in the heat map at the right-side panel (red indicates increase and green indicate lower than PC) **(D)** Representative confocal images of the indicated TNBC PC and their Paclitaxel (PDTP-P) and Doxorubicin (PDTP-D) PDTP stained with EGFR and DAPI as described in Materials and Methods. Scale bar, 10 µm. **(E)** Mean fluorescence intensity of total cellular EGFR and cell-membrane localized EGFR were quantitated using Image J software in PC, PDTP-P and PDTP-D of MDA-MB468. Similarly, membrane and cytoplasmic EGFR were quantitated in PC of MDA-MB468 and HS578T **(F)** Effects of short-term treatment of Paclitaxel (10 nM) on the levels of EGFR and p-EGFR protein in the indicated cell lines as estimated by WB analysis at different time points of treatment. Heatmaps presented in the right panel show the fold change in the protein expression quantified by densitometric analysis using ImageJ and expressed as folds of untreated (UT) control cells **(G)** The indicated TNBC cell lines were treated with different concentrations of paclitaxel (see Materials and Methods) for 24 hours and levels EGFR and pEGFR proteins were assessed by WB. Beta-tubulin served as their loading control. Heatmaps at the right panel show the fold change in the protein expression quantified by densitometric analysis using ImageJ and expressed as folds of untreated (UT) control of respective cells.

To understand the molecular reprogramming associated with the PDTP state in TNBC cells, we performed proteomic analysis. We subjected the total cell lysates of MDA-MB-468 parental cell (PC) and PDTP-Paclitaxel (PDTP-P) cells to Mass spectrometry analysis as described under the material and method section. The analysis identified 207 distinct proteins and out of these 67 proteins were differentially expressed in a significant manner between PDTP-P and PC. We further filtered out the proteins with more than 1.5 folds difference in the expression, and with this MDA-MB-468 PDTP-P showed 13 proteins that were upregulated and 46 proteins that were downregulated compared to PC (Supplementary Table 1). As shown in the volcano plot of the differentially expressed proteins (Fig. 1B) identified by the mass spectrometric analysis, EGFR was found to be the most significantly downregulated protein (p-value =2.15963E-05) in PDTP-P. In a parallel approach to characterize the phospho-proteomic changes in PDTPs we have probed the total cellular lysates from the PC and PDTPs with anti-phospho-tyrosine antibody to analyze the gross differential signals between PCs and PDTPs. We observed a higher (upto 1.38 folds) anti-phospho-tyrosine signal in PDTPs (Fig.S1A). We then analyzed the expression and tyrosine phosphorylation of EGFR in order to understand the alterations in the activation status of EGFR using Western blotting (WB). Interestingly, we found that EGFR protein expression is drastically reduced in most of the PDTPs (Paclitaxel, PDTP-P; doxorubicin, PDTP-D; and cisplatin, PDTP-C) as compared to their respective PCs irrespective of the TNBC subtype or chemotherapy (Fig. 1C, Fig. S1B). Surprisingly, the tyrosine phosphorylation status of EGFR at the activation site Y^1068^ was up to 55 folds higher, while the pEGFR to EGFR ratios was elevated up to 125 folds in PDTPs (Fig. 1C, Fig. S1B). We also performed immunofluorescence (IF) staining of different PCs and PDTPs lines to check the sub-cellular localization of EGFR in steady-state conditions. We have found that EGFR is highly expressed in parental cells and primarily localized at the cellular surface; however, EGFR had a consistently lower expression in PDTP-P and PDTP-D cells and was predominantly localized in the perinuclear regions in vesicular structures (Fig. 1D). The quantification of IF staining at the total cellular and cell membrane level also suggest that cell surface localization of EGFR is significantly lower in PDTPs while TNBC PCs predominantly express EGFR at cell-surface (Fig. 1E) as compared to cytoplasm. To understand how chemotherapy affects EGFR protein expression and activation in PC TNBC cells, we checked dose- and time-dependent expression and phosphorylation patterns of EGFR in the MDA-MB-468 cell line using WB analysis. As seen in Figures 1F and G, paclitaxel treatments in PC did not affect the total EGFR levels up to 48 h of treatment, even at different dosages. These results indicate that changes in EGFR phosphorylation and downregulation are not early events in TNBC cells following chemotherapy, which suggests that possibly the downregulation of the receptor may have commenced during the selection of PDTPs with a hyper-phosphorylated status.

### A Low EGFR expression state is associated with poor survival in chemotherapy-treated breast cancer patients and is associated with low pCR and recurrence-free survival in TNBC patients

As we have observed a dramatic downregulation of EGFR in TNBC PDTPs, we wanted to assess the clinical relevance of our findings in breast cancer patients using publicly available TCGA data. First, we analyzed the gene expression of EGFR in human TNBC samples from different molecular subtypes (Fig. 2A). It was observed that EGFR expression is highest in BL1 and BL2 subtypes of TNBC tumors. Further, we have analyzed the dependency of breast cancer cell lines on EGFR expression using Dependency Map (DepMap) at the https://ualcan.path.uab.edu/ platform. Normalized EGFR gene effect scores derived from CRISPR knockout screens (inferenced as Chronos) were downloaded and sorted from the highest affected cell line to the lowest or non-affected ones. As seen in Figure 2B, most of the breast cancer cell lines have negative Chronos scores that indicate proliferation inhibition and/or death following EGFR knockout suggesting an important role of EGFR-mediated cell signaling in the survival of breast cancer cells. This data implies that the downregulation of EGFR at the protein level might be compensated by the hyper-activation of EGFR in TNBC PDTPs to support the proliferation (Fig. 1C, Fig. 1SB). Next, we analyzed the association of EGFR gene expression on the recurrence-free survival (RFS) and distant-metastasis-free survival (DMFS) of TNBC patients with different levels of disease progression under systemic chemotherapy. Kaplan-Meier survival plots show that in chemotherapy-treated TNBC patients, low EGFR expression is an indicator of very poor RFS (HR=0.34, P<0.05) (Fig. 2C, D) as well as DMFS (Fig. 2E) in high-grade lymph-node positive TNBC patients (HR=0.13, p value=0.0037). Further, we have also examined the association of EGFR protein expression in the survival of breast cancer patients using proteomics data available at the TCPA (The Cancer Protein Atlas) from the TCGA-BRCA cohort (901 patients) and found a significant correlation between lower EGFR protein with poor survival (Fig. 2F). These results support the finding that lower expression of EGFR protein in breast cancer is a significant predictor of poor survival. On the other hand, EGFR expression in tumors has no prognostic value in untreated breast cancer (Fig. S2A) or TNBC patients (Fig. S2B). Interestingly, analysis of EGFR expression was also found to be lower in metastatic breast tumors compared to primary tumors (Fig. S2C). As we have found that in PDTPs EGFR is very significantly phosphorylated at the Y^1068^, we also analyzed the proteins that are highly correlated with EGFR Y^1068^ expression using TCPA cohort. It was seen that pEGFR is strongly correlated with several other oncogenic phosphoproteins like SHP2, HER2 and SRC as well as activated RTK signaling like PI3K and MAPK (Fig. S2D). Further, we analyzed a breast cancer gene expression array dataset GSE22513 from breast biopsy of patient groups gaining pCR or non-pCR with Taxane using the GEOR platform. We found that patients attaining pCR with Taxane had a significantly higher expression of EGFR (Log Fold Change, 2.2639385, p value=1.60E-03) compared to the cohort that has non-pCR (Fig. 2G). In light of the above findings, we examined whether EGFR gene expression is linked to response to chemotherapy and can be a predictive biomarker for pCR and RFS in breast cancer patients. For that, we have performed a ROC (receiver operating characteristic) analysis with the ROC plotter to compute pathological complete response (pCR) and RFS in chemotherapy-treated TNBC patients based on AUC (Area Under the Curve) values (Fekete and Győrffy 2019). As seen in Figure 2F, interestingly, in TNBC patients, EGFR expression is a very strong predictor of RFS in chemotherapy-treated metastatic patients with an AUC value of 0.813 (Fig. 2H). TNBC responders show a very high expression of EGFR as compared to non-responses (Fig. 2D). This patient data analysis suggests that EGFR expression is a robust predictor of chemotherapy response, especially in TNBC patients, which is in line with our results (Fig. 1, S1), where we observed a low expression of EGFR in PDTPs from different TNBC subtypes.

**Figure 2.**
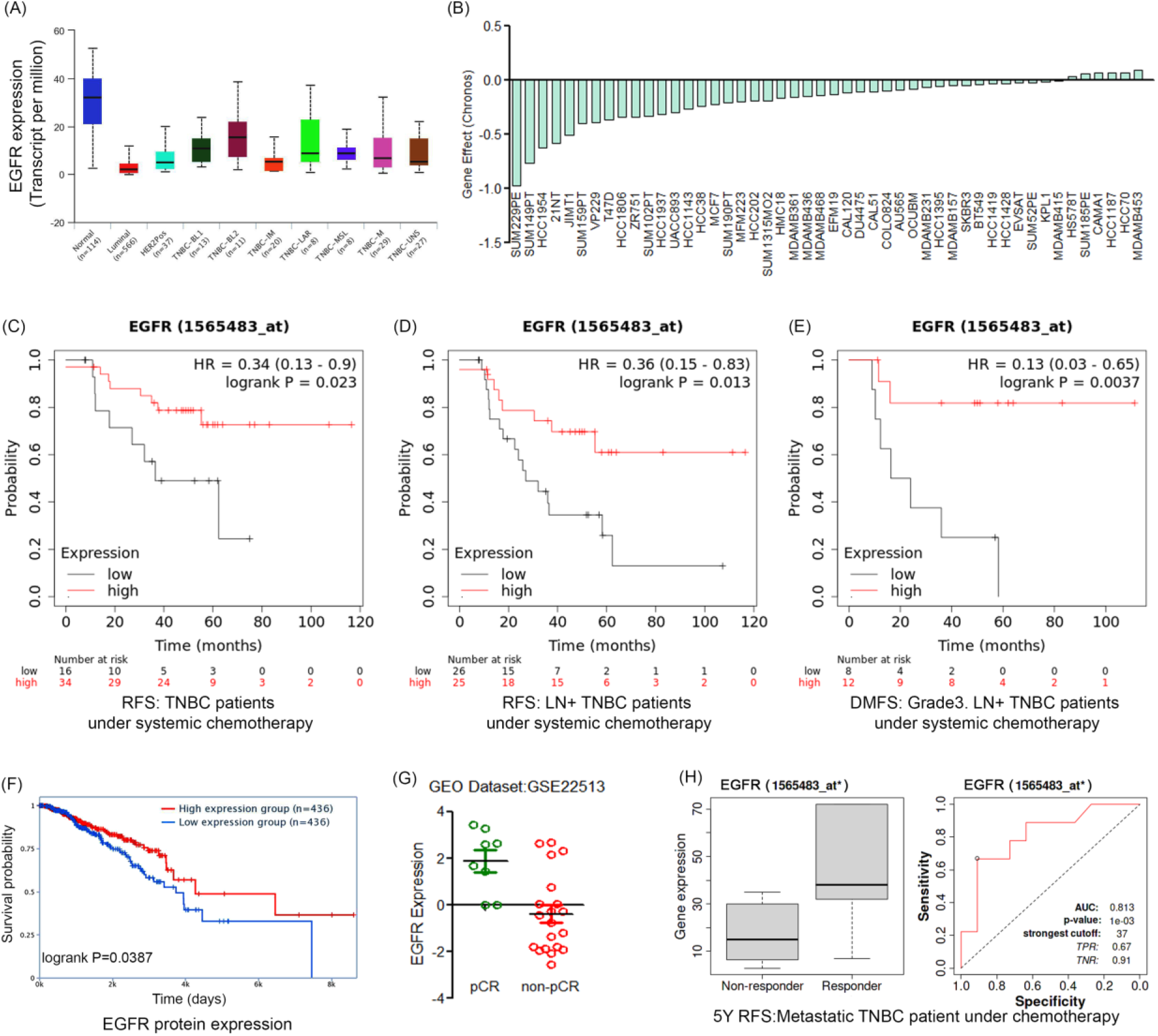
Low EGFR expression is a strong indicator of poor outcomes in chemotherapy-treated TNBC patients. **(A)** EGFR mRNA expression (transcript per million) in human normal breast tissues and in TNBC patient tissues belonging to different TNBC subtypes are shown as a box plot derived from TCGA data. **(B)** Waterfall plot showing the effect of EGFR gene CRISPR gene knockout in breast cancer cell lines in the order of intensity represented as Chronos scores. **(C)** Kaplan-Meier plot showing RFS in TNBC patients under chemotherapy. Similarly, **(D)** RFS is shown using mRNA expressions of EGFR in lymph node-positive (LN+) TNBC patient samples treated with chemotherapy and stratified into a high or low expression using the auto expression value as the cut-off point. **(E)** DMFS in high-grade LN+ TNBC patients under chemotherapy is shown in Kaplan-Meier plot. Expression of EGFR in TNBC tissues was stratified into a high or low expression using auto-cut values. Gene expression of all the breast cancer patients having low (Black) or high (Red) EGFR expression is defined by their median values. The HR score and corresponding p-value for the Log-rank test is indicated. **(F)** Kaplan-Meier plot showing breast cancer patients’ survival as a function of EGFR protein levels using the proteomic TCPA data. **(G)** Dot plot showing differential expression of EGFR gene in tumors of breast cancer patients achieving pCR and non-pCR as analysed from gene expression dataset (GSE22513) using GEOR on GEO in 28 patient samples treated with Taxane. **(H)** Box plot showing gene expression of EGFR in responders vs. non-responders and ROC curve comparing sensitivity and specificity of EGFR gene for classifying responders vs non-responders LN+ TNBC patients with chemotherapy

### NDRG1 upregulation mediates lysosomal degradation of EGFR in Proliferating drug-tolerant persister TNBC cells and predicts poor recurrence-free survival in chemotherapy-treated TNBC patients

As we have observed that EGFR protein levels are downregulated in TNBC PDTPs (Fig. 1C, S1B) together with the change in the subcellular localization (Fig. 1D, E), we sought to determine if decreased protein levels are due to lysosomal mediated degradation of the receptor. In order to investigate that, we performed IF of EGFR and lysosomal marker protein LAMP1 in PC and PDTPs in steady-state conditions and with the treatment of lysosomal inhibitor chloroquine (CQ). and we observed that EGFR is mostly localized at the cell membrane and shows a minimal co-localization with lysosomal marker LAMP1 (Fig. 3C). EGFR was seen to be significantly co-localized with LAMP1 in the MDA-MB-468 and HS578T PDTP-P (Fig. 3A-C), and also in the PDTP-D cells (Fig. S3A-E). Co-localization analysis of cells in TNBC PC and PDTPs suggests that EGFR is mostly located in a lysosomal compartment in chemotherapy-tolerant persister cells (Fig. 3C; Fig. S3B-E). Treatment with CQ in PDTPs resulted in the accumulation of EGFR in the lysosomal compartment and increased intensity of anti-EGFR staining in these cells, suggesting that EGFR is mostly degraded in PDTP cells. Further, we performed a WB analysis of CQ-treated MDA-MB-468 PC and PDTP-P to determine the recovered EGFR levels from PDTP cells’ lysosomal degradation. We have observed a substantial amount (7 folds of PC) of EGFR recovery under 12 h of CQ treatment in MDA-468-PDTP-P but not in MDA-468-PC cells (Fig. 3D), suggesting that EGFR is rapidly degraded in the proliferating drug-tolerant persister cells.

**Figure 3.**
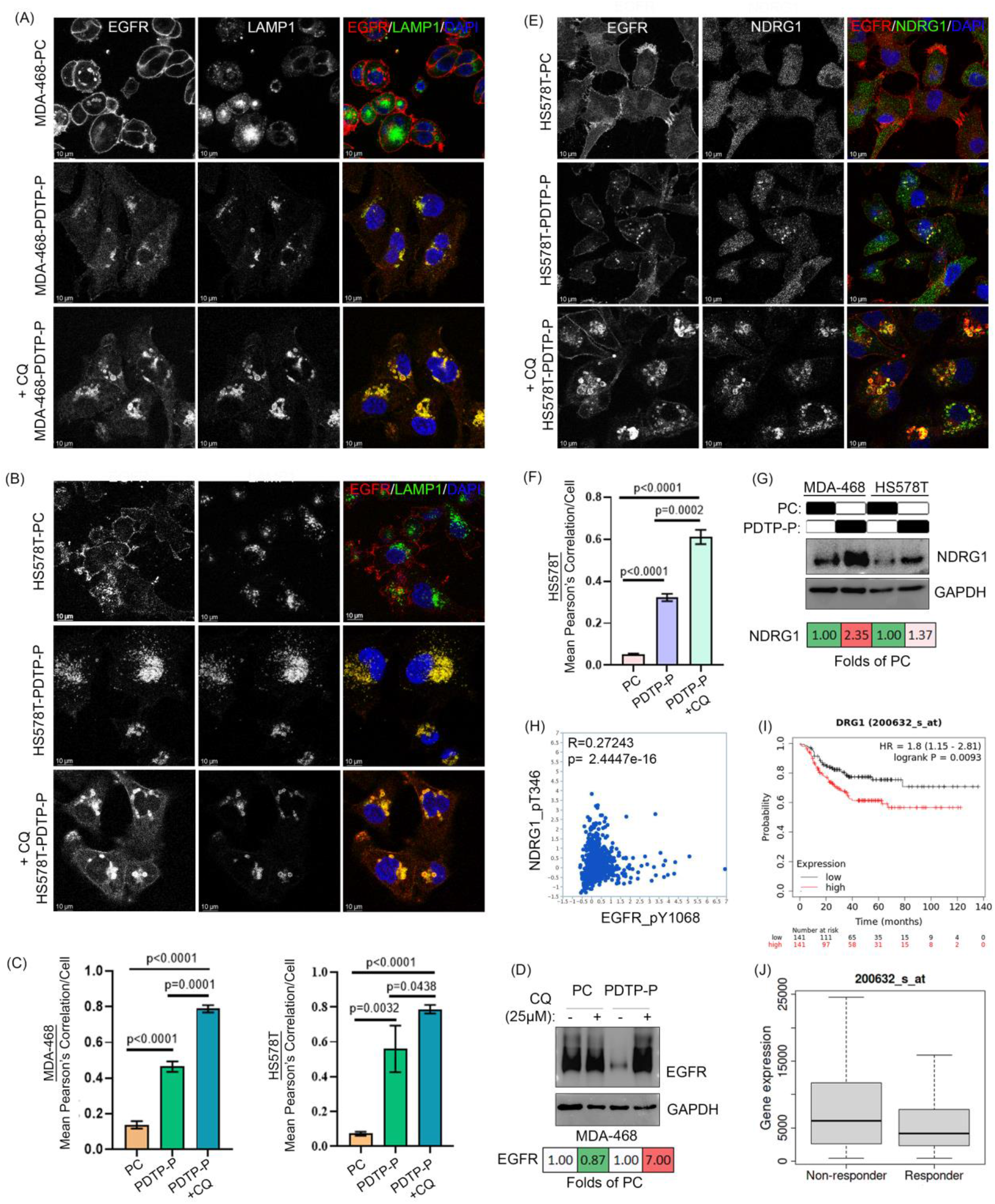
EGFR is trafficked to the lysosomal pathway for degradation through increased NDRG1 in TNBC PDTPs. **(A)** and **(B)** Representative confocal images of the indicated PC and their Paclitaxel PDTP TNBC cells stained with EGFR, LAMP1 and DAPI +/-treatment with CQ (25 µM for 12 hrs) as described in Materials and Methods. Scale bar, 10 µm. **(C)** Bar graph showing mean pearson’s correlation of EGFR and LAMP1 per cell depicting colocalization of these proteins were quantitated using LAS X software in PC, PDTP-P and PDTP-P+CQ of MDA-MB-468 and HS578T cells **(D)** Levels of EGFR determined in MDA-MB-468 PC and their Paclitaxel PDTPs with and without CQ (25 µM for 12 hrs) treatment as determined by WB analysis and quantified using densitometric analysis by ImageJ, presented as folds of PC. **(E)** Representative confocal images of the HS578T PC and their Paclitaxel PDTPs stained with EGFR, NDRG1 and DAPI +/-treatment with CQ (25 µM for 12 hrs) as described in Materials and Methods. Scale bar, 10 µm. (F) Bar graph showing mean pearson’s correlation of EGFR and NDRG1 per cell depicting colocalization of these proteins were quantitated using LAS X software in PC, PDTP-P and PDTP-P+CQ of HS578T cells. **(G)** Levels of NDRG1 were determined in PC MDA-MB-468 and HS578T and their Paclitaxel PDTP. GAPDH served as their loading control. The expression levels of the indicated proteins were assessed by WB, and band intensities were quantified by ImageJ software and presented as fold of control in the heatmaps (red indicates increase, and green indicates lower than PC). **(H)** Correlation plot showing the spearman’s correlation between pEGFR pY^1068^ and pNDRG1 p^T346^ protein expression in breast cancer using the BRCA cohort on TCPA. **(I)** mRNA expression of NDRG1 in TNBC chemotherapy-treated patient tissue samples was stratified into a high or low expression using the median expression value as the cut-off point. The corresponding p-value for the Log-rank test in chemotherapy-treated TNBC patient tissue samples was shown. Higher mRNA expressions of NDRG1 were significantly associated with poorer Recurrence-free survival (RFS) in TNBC chemotherapy-treated TNBC patients. Gene expression of all the TNBC chemo patient’s samples having low (Black) or high (Red) NDRG1 expression defined by their median value is shown. **(J)** Box plot of gene expression of NDRG1 in responders vs non-responders in chemotherapy-treatment TNBC patients cohort.

It is well known that NDRG1 is a unique regulator of multi-vesicular body formation dynamics and channelization trafficking of several cell surface receptors (including HER family of receptors) (Kovacevic et al. 2016; Verma et al. 2017) from endosome to lysosome. NDRG1 expression is also known to modulate multidrug resistance in several tumor types (Verma et al. 2017; Yang et al. 2021), therefore, we checked the expression of NDRG1 and its co-localization with EGFR. Using IF staining of NDRG1 and EGFR in HS578T-PC and -PDTP-P cells (Fig. 3E) we show an increased expression of NDRG1 in PDTP-P. It also co-localizes with EGFR in PDTP-P (Fig. 3E). Chloroquine treatment leads to accumulation of EGFR in lysosomal compartments where NDRG1 significantly co-localized with EGFR (Fig. 3F). We have also confirmed using WB analysis that NDRG1 was upregulated (Fig. 3G) in drug-tolerant proliferating persister TNBC cells compared to their PC counterparts. These results indicate that NDRG1 upregulation enhanced the lysosomal trafficking of EGFR in Proliferating drug-tolerant persister cells.

As we see a strong increase in NDRG1 levels in TNBC PDTP cells together with upregulation of the pEGFR signal, we checked the correlation of these two proteins in human breast cancer samples using the TCPA cohort. As seen in Figure 3H, pEGFR Y^1068^ significantly correlates with activated NDRG1 (Spearman’s correlation coefficient= 0.27243; p value=2.4447e-16). Furthermore, we sought to evaluate the role of NDRG1 in the RFS of chemotherapy-treated TNBC patients to understand its pathological relevance. Kaplan Meier plots from the TCGA cohort (Fig. 3I) shows that higher NDRG1 expression is associated with poor relapse-free survival in TNBC patient with chemotherapy. It was evident from the ROC curve (AUC value, 0.592, p=2e-02) (Fig. S3F) in TNBC patients who have better 5-year RFS with chemotherapy express less NDRG1 (Fig. 3J); therefore, NDRG1 expression levels can be used to predict the survival in TNBC patients under treatment. These findings correlate well with results seen in PDTP cells.

### EGFR-to-Src/Lyn/Fyn switching pivots a hyper-phosphorylated tyrosine kinase state in proliferating drug-tolerant persister TNBC cells and activates downstream RTK signaling

We have seen a drastic increase in EGFR phosphorylation in TNBC PDTPs; however, the turnover of EGFR is high due to the NDRG1-mediated lysosomal trafficking of the receptor. With these observations, we checked the effect of the EGFR hyper-phosphorylation on the downstream RTK signaling using WB analysis. We have observed that MDA-468-PDTP-P and HS568-PDTP-P cells had a substantial upregulation of pSTAT3, MAP kinases (pERK1/2 and pP38), and pAKT (Fig. 4A) levels which suggest the robust RTK signalling. However, it was surprising that despite considerably low EGFR expression due to NDRG1 mediated degradation, RTK downstream signaling is robustly upregulated in the TNBC PDTPs. We have seen a strong correlation of pEGFR with pSRC (Fig. S2D), and it is known that EGFR downstream signals can be sustained by its downstream cytosolic kinases from the Src family (Canonici et al. 2020; Formisano et al. 2014) as EGFR directly activates them. Therefore, we probed the expression and activation status of Src family kinases’ in TNBC PDTPs. Very interestingly, we observed that TNBC PDTPs have a strong upregulation of both phospho as well as the total level of Src (up to 55 folds of PC), together with a huge upregulation of Lyn and Fyn kinases at the total protein levels (Fig. 4B, Fig. S4A). These results indicate a molecular switching of EGFR to the Src family to keep the elevated downstream RTK signaling in the PDTPs that compensates for the EGFR downregulation and produces a hyper-phosphorylation state of the EGFR-Src kinases axis in TNBC PDTP cells.

**Figure 4.**
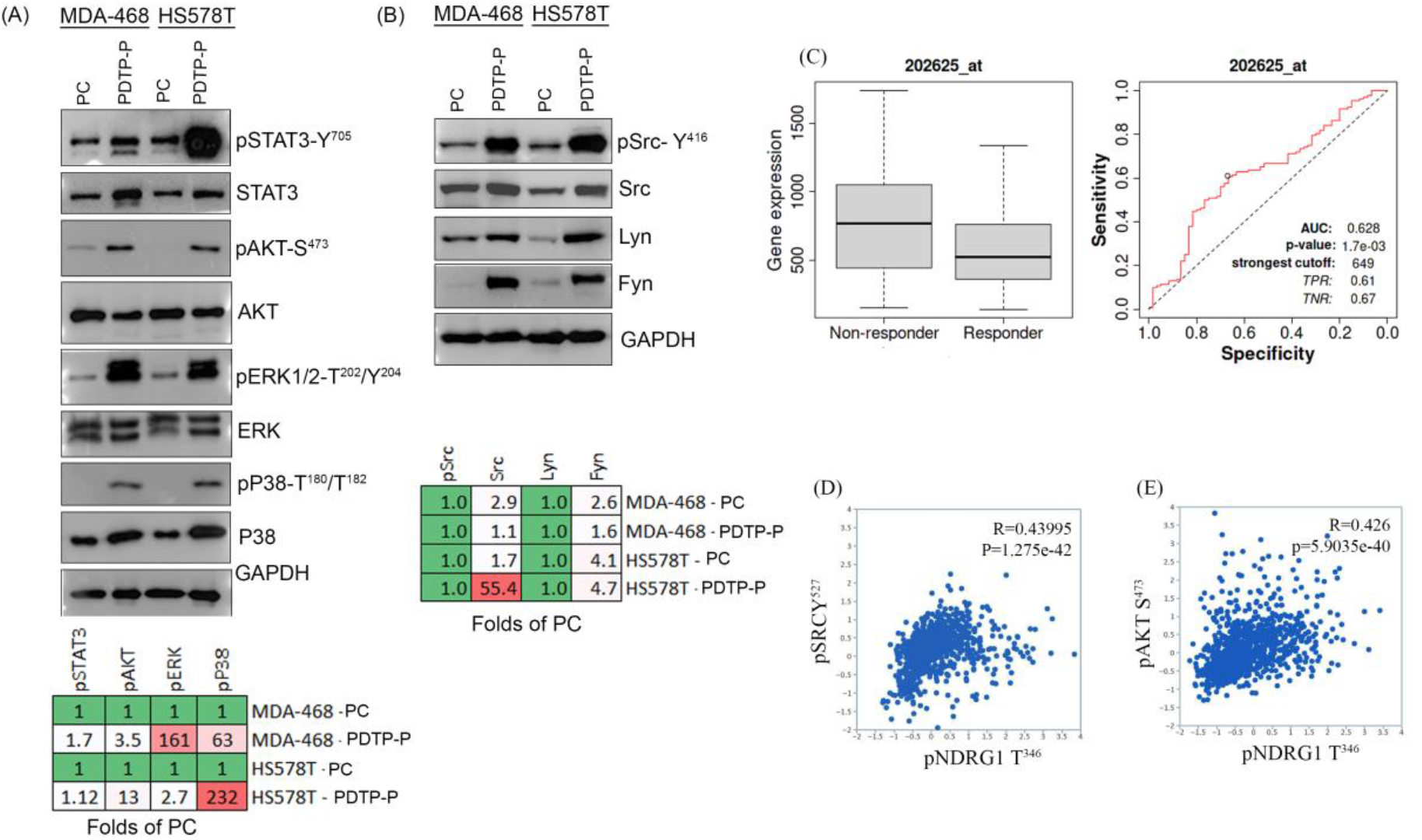
TNBC PDTPs have elevated activation of RTK signaling due to increased expression of Src family tyrosine kinases which indicate non-response in TNBC patients with chemotherapy and positive association with NDRG1. **(A)** Levels of STAT3, AKT, ERK, and p38MAPK and their phosphorylated forms were determined in MDA-MB-468 and HS578T PC and their Paclitaxel PDTP cells. GAPDH served as their loading control. The expression levels of the indicated proteins were assessed by WB, and band intensities were quantified by ImageJ software and presented as fold of PC in the heatmaps (red indicates increase, and green indicates lower than PC). **(B)** Levels of Src, p-Src, Lyn, and Fyn were determined in PC MDA-MB-468 and HS578T TNBC cell line and their Paclitaxel PDTP cells. GAPDH served as the loading control. The expression levels of the indicated proteins were assessed by WB, and band intensities were quantified by ImageJ software and presented as fold of control in the heatmap. **(C)** Box plot of gene expression of Lyn in responders vs non responders to chemotherapy treatment in TNBC patient samples. ROC curve comparing sensitivity and specificity of Lyn gene for classifying responders vs non responders to the chemo-treatment in TNBC patient samples. **(D)** Spearman’s Correlation plots between p-SRC and p-NDRG1 and **(E)** between p-Akt and p-NDRG1 in breast cancer samples obtained from BRCA TCPA datasets.

Src family kinases are well implicated in different tumors for poor prognosis in patients; we have analyzed the association of Lyn kinase with the pCR of TNBC patients with chemotherapy. As seen in Figure 4C, ROC curve and the expression of Lyn kinase in TNBC patients, it was observed that Lyn is significantly upregulated in TNBC patients who do not achieve pCR with chemotherapy, with a significant (p=1.73^-03^) AUC value of 0.628. This result indicates a strong predictive value that TNBC patients with high Lyn levels end up with residual disease, which is in line with our findings in TNBC PDTPs (Fig. 4B and Fig. S4). Considering these results, we have also analyzed the gene expression status of the tri-gene signature (SRC-LYN-FYN) in breast cancer patients, and we found this gene signature is significantly high in metastatic breast cancer (Fig. S4B) compared to the primary tumor.

In addition, we were curious to understand if NDRG1 protein correlates with pSrc and RTK downstream signals in breast cancer. To find this answer, we did a correlation analysis between activated states of pSrc-Y^527^ and pNDRG1-T^346^ in breast cancer patients using publicly available phospho-proteomic data from TCPA in the TCGA-BRCA cohort. Surprisingly, we found the activated Src protein levels highly correlated (spearman’s correlation, R=0.439, p=1.275e-42) with the pNDRG1 levels (Fig. 4D) in the BRCA patient cohort indicating a positive correlation between these phospho-proteins in breast cancer. Similarly, pAKT-S^473^ also positively correlated with pNDRG1 levels (spearman’s correlation, R=0.426, p=5.9035e-40) in breast cancer patients (Fig. 4E). The gene expression and the phospho-proteomic data from TCGA-BRCA patient cohort as well as our findings in Fig. 3E-J and Fig. 4B, suggest a positive co-activation effect of these proteins on each other in breast cancer.

### Paclitaxel-tolerant proliferating persister TNBC cells show higher sensitivity to EGFR tyrosine kinase inhibitor

Significantly lower levels of EGFR and hyper-phosphorylation status of EGFR, in PDTPs we hypothesized that these cells can be more susceptible and effectively targeted using EGFR-directed tyrosine kinase inhibitors. We then tested the response of Gefitinib, a well-documented targeted therapy against EGFR in epithelial tumors, especially in lung and breast cancers (Le and Gerber 2019). Dose-response curves of Gefitinib in PC and PDTP-P in different TNBC lines (Fig. 5A) show that paclitaxel tolerant proliferating persister TNBC cells are more sensitive to Gefitinib induced cell death as assessed by MTT cell viability assays. We also tested the potential of Neratinib, a newer generation of TKI that inhibits both ligand-dependent and independent phosphorylated HER2 and EGFR and is a potential therapeutic agent in breast cancer treatment and we observed that. that HS578T-PDTP-P is also more sensitive to Neratinib (Figure S5).

**Figure 5.**
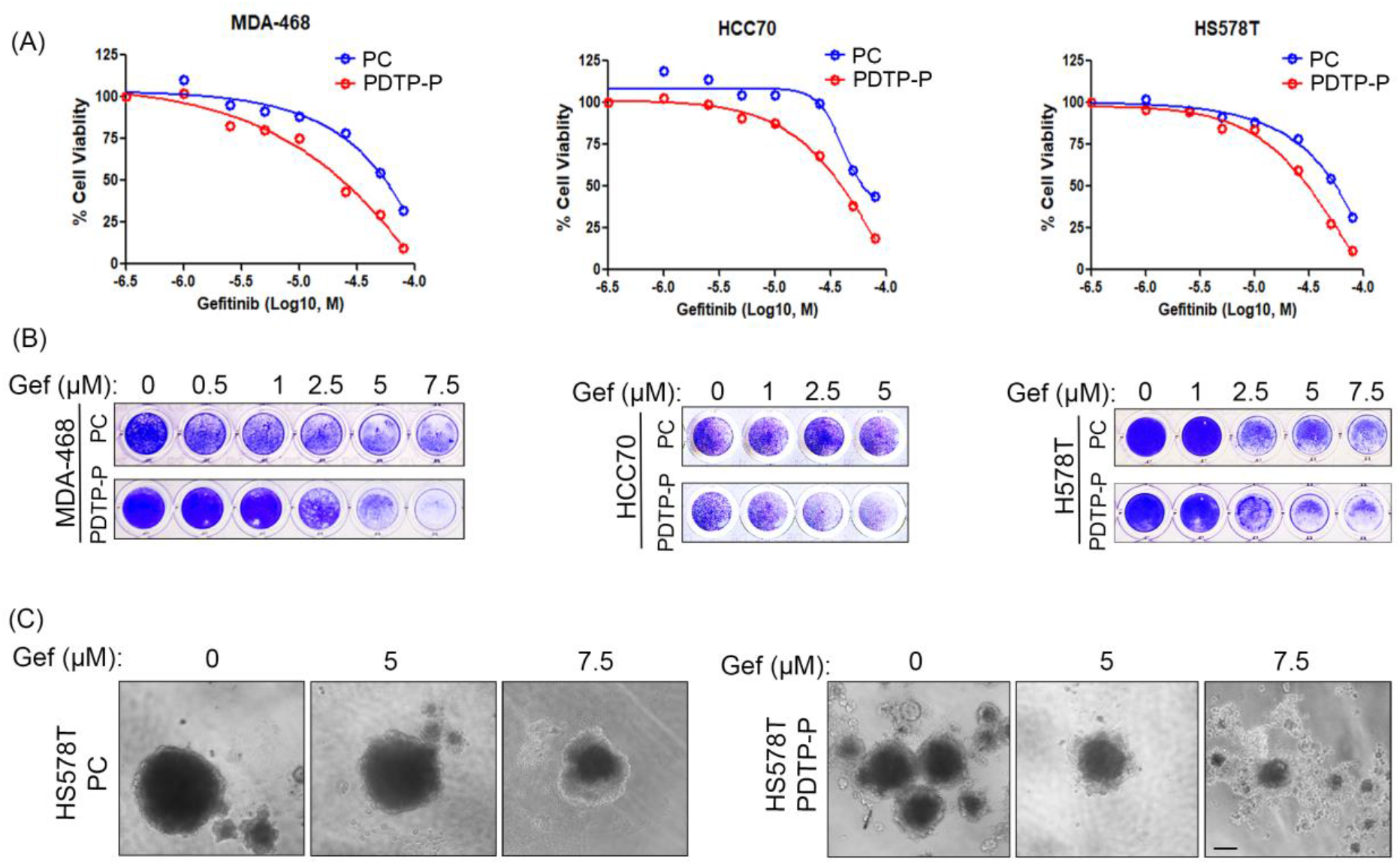
TNBC PDTPs are more sensitive to EGFR inhibitor Gefitinib. **(A)** Dose-response curves of varying concentrations of Gefitinib in the indicated cell lines (PDTP and PDTP-P) treated for 72 hours. Dose-response curves are presented as means of three replicates. **(B)** Effects of Gefitinib on cell viability. The indicated PC, TNBC cell lines and their Paclitaxel PDTP were treated with the indicated doses of the drugs for 72 hours and stained with crystal violet. Representative pictures of reproducible effects from two to three independent experiments are shown. **(C)** Effects of drug combinations on HS578T-PC and PDTP-P spheroid growth with and without Gefitinib (0, 5 and 7.5 µM) treatment. Representative micrographs (10x magnification) images of day 15 old spheroids (n = 6 spheroids) are shown. Control untreated and Gefitinib-treated spheroids with a single agent with the indicated drug levels are shown. Scale bar, 100 µm.

Similarly, the colony formation assay (Fig. 5B) indicates that PDTP-P cells lose their potential to grow and form viable colonies in lower doses of Gefitinib compared to their PC counterparts in different subtypes of TNBC. Collectively, these results show that due to a higher dependency on EGFR hyper-phosphorylated state, the PDTP-P cells are more vulnerable to EGFR TKI-induced killing.

Further, we analyzed the Gefitinib susceptibility of PC and PDTP-P in 3D spheroids in HS578T as they are enriched in stem cells and have a greater capacity to generate 3D spheroids than other TNBC MSL cell types. We observed that HS578T-PDTP-P spheroids are highly susceptible to Gefitinib as compared to the parental cells (Fig. 5C). These results further strengthen the above finding in 2D cultures and strongly suggest the potential of EGFR TKI to induce cell death more selectively in drug-tolerant proliferating cells and kill tumors cells in 3D growth where stemness features of cancer cells are more enriched.

### Gefitinib re-sensitizes proliferating drug-tolerant persister TNBC cells to paclitaxel by strongly hampering EGFR-mediated Src activation and downstream RTK signaling

In light of the above exciting findings that demonstrate a higher vulnerability of TNBC PDTPs to EGFR TKIs, an effect independent of TNBC subtypes, we went ahead and tested the response of paclitaxel and Gefitinib combination therapies in paclitaxel PDTP TNBC lines. Remarkably, the combination of paclitaxel and Gefitinib resulted in a much higher percentage of reduction in the colony formation capacities of PDTP-P TNBC cells as compared to the PCs in all tested lines (Fig. 6A). It is interesting to see that the two drugs in combination were working synergistically while there is a minimal or less profound effect on the viability of cells when treated with paclitaxel or EGFR TKI alone in both PC and PDTP-P cells.

**Figure 6.**
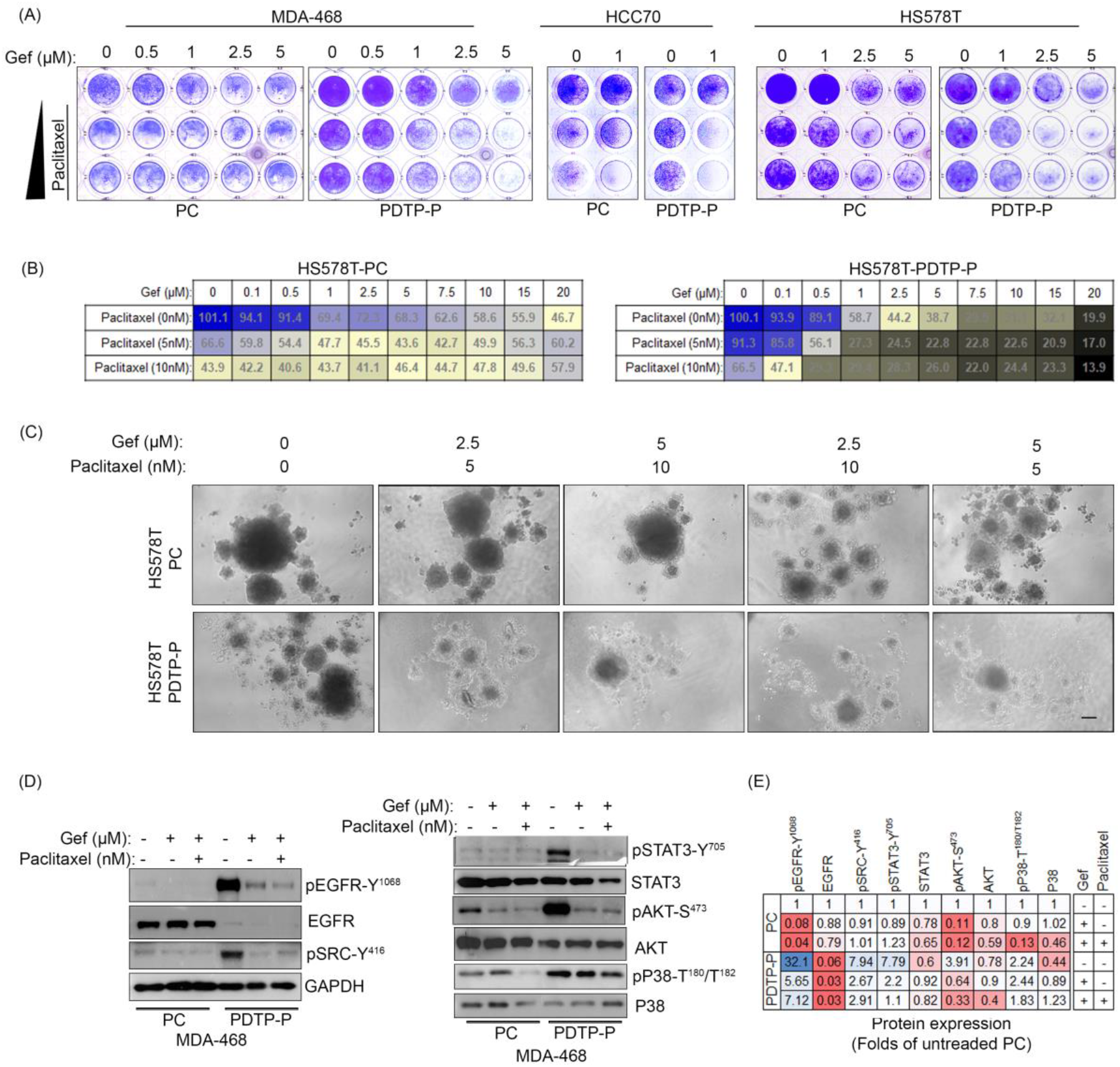
Combination of Gefitinib and paclitaxel selectively induce cell death in PDTP-P cells by suppressing EGFR-Src hyper-phosphorylation and downregulation of Lyn and Fyn kinases. **(A)** Effects of Gefitinib and Paclitaxel on cell viability. The indicated PC TNBC cell lines and their paclitaxel (PDTP-P) were treated with the indicated doses of the Gefitinib and Paclitaxel drugs for 72 hours and stained with crystal violet. Representative pictures of reproducible effects from two to three independent experiments are shown. **(B)** The indicated cell lines were treated with Gefitinib and Paclitaxel at the indicated dose (see Materials and Methods). Cell viability (MTT assay) is presented as the percentage viability of untreated cells. The mean values of three experiments are shown. **(C)** Effects of drug combinations on PC- and PDTP-P-HS578T-spheroid growth. Representative micrographs (10x magnification) images of day 15 day-old spheroids are shown. Control, untreated, and Gefitinib treated spheroids with a single agent and with the Paclitaxel drug levels are shown. Scale bar, 100 µm. **(D)** The expression levels of the indicated proteins were assessed in MDA-MB-468 cell line treated with Gefitinib and Paclitaxel as indicated by WB. **(E)** Heatmaps showing fold changes in MDA-MB-468-PDTP-P and HS56T-PDTP-P as compared to corresponding parental cells are shown. The expression of Lyn and Fyn kinases Quantitative band intensities were quantified by ImageJ software and presented as fold of control in the heatmaps.

Further, we performed a cell viability assay using MTT to see the quantitative response of Gefitinib and paclitaxel alone and in combination in HS578T-PC and –PDTP-P cells (Fig. 6B). We observed a very effective combinatorial effect of these two drugs in PDTP-P cells starting from a Gefitinib concentration as low as

0.5 µM with 5 and 10nM of paclitaxel, where PDTP-P cells are resistant to these paclitaxel concentrations alone, whereas, these concentrations are around IC_50_ in the PCs and kill parental cells.

The effect of the combination of Gefitinib and paclitaxel was also tested on 3D spheroids generated from HS578-PC and –PDTP-P. We see a strong dose-dependent growth inhibitory effect of this combination therapy in HS578T-PC, while a cytotoxic effect is seen in HS578T-PDTP-P spheroids, even starting from the minimal concentrations tested (Fig. 6C). These results collectively indicate a very strong effect of Gefitinib when combined with paclitaxel in PDTP-P cells, while a modest growth inhibitory effect was seen only in 3D cultures in HS578T-PC.

Lastly, we sought to investigate how the combination therapy of EGFR TKI with paclitaxel predominantly induces cell death in TNBC-PDTP-P cells. To answer this question, we have again extensively examined the robust hyper-phosphorylated EGFR-Src signaling hub and the RTK downstream signaling in the drug combination using WB analyses of these phospho and total proteins. We have discovered that Gefitinib by itself dramatically reduced the hyper-phosphorylated levels of EGFR and Src in MDA-468-PDTP-P to relative levels present in parental cells (Fig. 6D) while also effected the activation of STAT3 and AKT in these cells. Interestingly, the combination of Gefitinib with paclitaxel reduced the total level of Lyn and Fyn kinases in MDA-MB-PDTP-P and HS578T-PC cells (Fig. S6A-D). These results indicate that while EGFR inhibitor suppressed pSrc and downstream RTK signaling, there is a strong downregulation of Src family kinases as an effect of combination therapy that disintegrates the tyrosine kinase hub built in the paclitaxel-tolerant TNBC cells supporting their survival and proliferation.

Altogether, these results suggest that the combination of EGFR TKI with paclitaxel can selectively eliminate the proliferating drug-tolerant persister cells that emerged post-chemotherapy due to the integration of EGFR-Src family kinases and building a robust hyper-phosphorylation status in the TNBC PDTPs (Fig. 7). Therefore, it could be beneficial to combine EGFR TKI to TNBC tumors that did not attain pCR; however, in these cases, the evolution of pEGFR, pSRC, Lyn, and NDRG1 expression can be determinant to assess the success of therapy.

**Figure 7.**
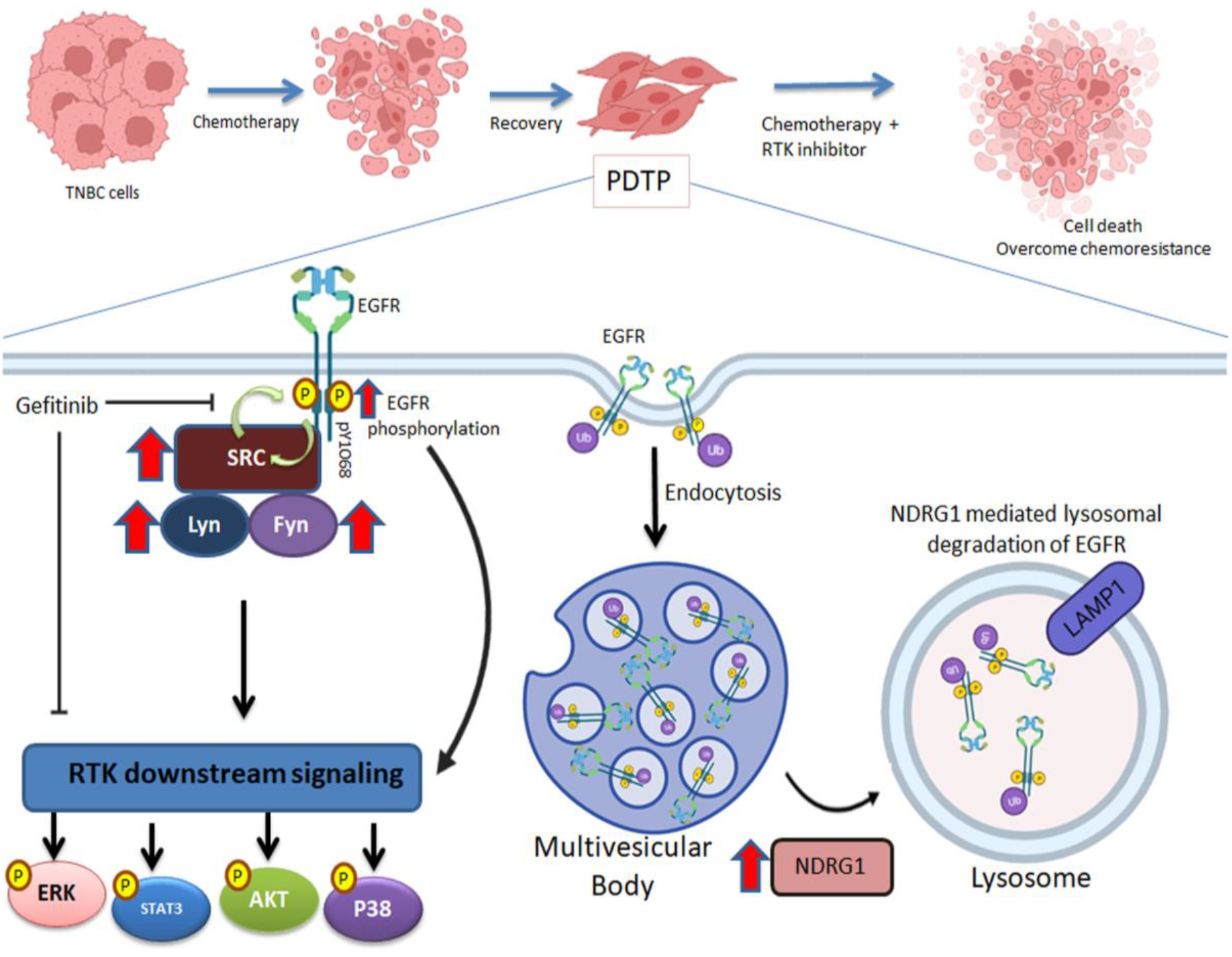
Schematic model summarizing the main findings of the study. TNBC cells treated with chemotherapeutic drugs will develop PDTP cells which have enhanced lysosomal trafficking of EGFR which is mediated by increased levels of NDRG1 to degrade EGFR. Despite having lower levels of total EGFR, PDTP cells have higher levels of Src family Kinase including Lyn and Fyn which maintains the activation of EGFR and leads to robust EGFR-Src tyrosine kinase axis. This EGFR-Src signalling axis is required for the downstream over-activation of STAT3, AKT and MAP kinases. Inhibiting EGFR-Src signaling axis with gefitinib and its combination with paclitaxel increases cell death in PDTP and thus depicting the unique switching of EGFR–Src family tyrosine kinases creating a vulnerability to EGFR TKI.

## Discussion

TNBC is one of the most aggressive breast cancer subtypes and has no clinically approved first-line targeted therapy. Therefore, TNBC patients are mainly treated with cytotoxic chemotherapies. The response rate of TNBC patients to chemotherapy is not more than 50% worldwide. The rest of the patients have drug-resistant residual disease and poor 5-year survival rates.

EGFR is one of the RTK whose overexpression has been considered one of the molecular hallmarks of TNBC (Park et al. 2014). In TNBC tumors, EGFR amplification is frequently seen and correlates with overexpressed protein levels, however, mutations are rarely seen. EGFR is strongly expressed in TNBC tumors (up to 67%), especially in Basal-like subtypes (Verma et al. 2017), and can be utilized for targeting TNBC cells where it is amplified and in combination with other therapies (Canonici et al. 2020; Sabatier et al. 2019; Savage et al. 2017). It has been shown that EGFR expression is associated with ABCG2 expression and function and ABCG2-mediated chemoresistance in TNBC cells. (Nedeljković and Damjanović 2019). In these cases, it has been demonstrated that EGFR inhibitors could reverse the ABCG2-mediated chemoresistance in both in-vitro and in-xenograft models.

Apart from a few studies, the role of EGFR in modulating response to chemotherapy in TNBC tumors has not been explored enough (You et al. 2021), especially in drug-tolerant diseases. Clinical studies indicate EGFR expression levels in primary TNBC tumors before chemotherapy can determine response to therapy (Tanei et al. 2016; Tang et al. 2012). Using clinical TNBC datasets, we have shown that EGFR expression is significantly lower in chemotherapy non-responder patients, and a higher level of EGFR expression correlates with better RFS in chemotherapy-treated TNBC patients. Tang et al. showed that overexpression of EGFR was significantly associated with pCR rate in patients with TNBC with neoadjuvant chemotherapy. However, most of these reports are based on total levels of EGFR, and the activation status of EGFR was not determined. Previous studies have evidenced that EGFR targeting in combination with cytosolic tyrosine kinases like Src and FAK family kinases can effectively suppress growth in different TNBC cells in vitro and in vivo animal models (Canonici et al. 2020; Verma et al. 2017); however, the association of EGFR with these cytosolic tyrosine kinases in chemotherapeutic response was not well explored in TNBC tumors cells.

In this study, we have shown that chemotherapy-tolerant cells from different TNBC subtypes undergo downregulation of EGFR levels due to NDRG-1 mediated lysosomal degradation (Menezes et al. 2017). However, the EGFR receptor has a huge increase in the activated EGFR state, with a concomitant upregulation of Src family kinases (Src, Lyn, and Fyn) which reflects a tyrosine kinase switching mechanism. We also show that higher NDRG1 expression in chemotherapy-treated TNBC tumors indicates an incomplete response to chemotherapy. Previous studies have suggested that EGFR levels regulate multidrug resistance in luminal breast cancer cells (Xu et al. 2011). In lung cancer where EGFR is highly repressed, it has been shown that chemotherapy renders lung cancer cells tolerant to EGFR inhibition via increased AXL-signaling (Aldonza et al. 2021). No previous study, however, clearly indicated the role of pEGFR status in chemotherapy-tolerant breast cancer cells.

It has been reported that stimulated EGFR alters molecular signaling in breast cancer cells and can induce metabolic shifts like increased glucose flux through the PI3K pathway, rendering these cells to metabolic targeting (Jung et al. 2019). We have shown that in TNBC PDTPs tyrosine kinases Src and EGFR were maintained in a hyper-phosphorylated state that sustained an over-activated RTK downstream signaling. This hyper-phosphorylation state makes these drug-tolerant cells molecularly distinct from parental TNBC cells. In TNBC PDTPs, although the hyper-phosphorylated EGFR-Src hub together with upregulated Lyn and Fyn kinases seems to provide the PDTPs a survival advantage in response to chemotherapy, at the same time, this tyrosine kinase axis creates a vulnerability to EGFR tyrosine kinase inhibitors. It has been shown in lung adenocarcinoma cells that Lyn can regulate EGFR phosphorylation levels ad siRNA-mediated knockdown of Lyn reduced activation of EGFR (Sutton et al. 2013). Therefore, it could be speculated that upregulated Lyn and Fyn kinases contribute to constitutive EGFR phosphorylation in TNBC PDTPs.

Recently, it has been shown that co-targeting of Src and EGFR can be an effective treatment for TNBC (Canonici et al. 2020). Src is known to sustain bypassed signals from EGFR and HER2 receptors in breast cancer cells in response to TKI treatment (Formisano et al. 2014; Rexer et al. 2011). Moreover, an interesting study using a kinome-wide siRNA screening identified the role of Src in chemoresistance in TNBC cells. These studies further substantiate our finding that Src activation is an active contributor to the survival of chemotherapy-tolerant cells.

We have exploited this hyper-phosphorylation state of TNBC PDTPs for selective targeting of chemo-tolerant cells and shown that EGFR TKI in combination with paclitaxel can be utilized to induce forceful cell death in post-chemotherapy tolerant TNBC cells. This combination treatment not just attacks the phosphorylation of Src and EGFR but also downregulates the protein levels of Lyn and Fyn kinases. Lyn is shown to be overexpressed in TNBC and is known to derive aggressive behavior in TNBC cells by integrating upstream RTK cellular signals such as from the c-KIT receptor (Tornillo et al. 2018).

In conclusion, our results identify a novel tyrosine kinase switching between EGFR-Src in chemotherapy-tolerant TNBC cell lines, a phenomenon that is independent of the TNBC molecular subtype. Further, the hyper-phosphorylation state of TNBC PDTPs built due to overexpression of Lyn and Fyn kinases supports the survival of these cells. Therefore, we highlight a rationale for combining EGFR TKI with chemotherapy agents like paclitaxel that can target TNBC drug-tolerant cells and provide a complete response by eliminating persister tumor cells post-chemotherapy.

## Materials and Methods

### Chemicals and Drugs

Chemotherapeutic agents used in the study were purchased from Sigma (Paclitaxel, catalog no. T1912; Doxorubicin, Catalog no. D1515, and Cisplatin, catalog no. 479306). EGFR inhibitors Gefitinib (catalog no. A10422) and Erlotinib (catalog no. A11416) were procured from Adooq Biosciences, USA. Thiazolyl Blue Tetrazolium Bromide (MTT) (M5655) and Chloroquine (catalog no. C6628) were procured from Sigma. Dithiothreitol (DTT, D9779), CaCl_2_(C1016) and Iodoacetamide(I-1149), Trypsin (T7575-1KT) were purchased from sigma. Urea (SRL 2113) was purchased from SRL.

### Antibodies

The following antibodies were purchased from Cell Signaling Technology (USA): rabbit anti-EGFR (4267), rabbit anti-phospho EGFR Y-1068 (3777), mouse anti–STAT3 (9139), and rabbit anti-phospho STAT3 Y-705 (9145), rabbit anti-Src (2123), rabbit anti-phospho Src Y-416 (6943), rabbit anti-Lyn (2796), rabbit anti-Fyn (4023), rabbit anti-AKT (9272), rabbit anti-phospho AKT S-473 (4058), rabbit anti-ERK1/2 (9102), rabbit anti-phospho ERK1/2 T-202 Y-204 (9101), rabbit anti-GAPDH (5174), rabbit anti-P38 (9212) rabbit anti-pP38 (4511), rabbit anti-NDRG1 (9485). Mouse anti–α-tubulin (T6074) from sigma Aldrich, mouse anti EGFR (ab30) abcam, and mouse anti LAMP1 (DSHB-H4A1). Alexa 488 (A11008, A11001) and 568 (A11004, A11011) were purchased from Invitrogen.

### Cell culture

Different subtypes of triple-negative breast cancer (TNBC) cell lines MDA-MB-468, MDA-MB-231, MDA-MB-453, HCC70, and HS578T, procured from ATCC (USA). These cells were maintained in RPMI 1460 medium (Gibco BRL, USA), in which l-Glutamine (2 mM) and sodium bicarbonate (2g/liter) were added. The media was supplemented with 10% fetal bovine serum (Gibco BRL, USA) and penicillin/ streptomycin (Invitrogen, USA). The Cells were cultured in a 5% CO_2_ humidified incubator at 37°C. Mycoplasma contamination was routinely checked with a commercially available kit (Biological Industries, Israel) as well as by DAPI staining.

### Mass spectrometric analysis

Mass spectrometry was performed for MDA MB 468 PC and MDA MB 468 PDTP-P cells, as per the previously described protocol (Narasimhan et al., 2019). Briefly, 1 × 10^6^ cells were suspended in 100 ul Laemmli buffer. The samples were boiled for 10 minutes at 95 °C, and then kept on ice for 5 minutes, followed by centrifugation at 13000x g for 15 minutes at 4°C, to prepare the whole cell lysate. The supernatant was then subjected to acetone precipitation by incubating with 1ml of ice-cold acetone for 1 hour at -20°C. Samples were then centrifuged at 13000x g for 15 minutes at 4°C.The protein pellet was air dried, and then resuspended in 6M urea for denaturation. Protein quantification was carried out using Bradford’s assay. The samples were then incubated at room temperature for 1 hour with 200mM DTT for reduction followed by 1 hour incubation with 200mM IAA for protein alkylation. 1mM Cacl2 was used to adjust the urea concentration to 0.6 M and the samples were then incubated with proteomic grade trypsin for 16 hours to carry out in-solution digestion. Peptides desalting was carried out using ZipTip U-C 18 spin columns (Cat. No. ZTC18M096, Millipore Corporation U.S.A.). The peptides were dried in a speed vac and then reconstituted with 0.1%Formic acid in water.

### SWATH analysis

LC-MS data acquisition on ESI-Q-TOF-MS (ABSCIEX, Triple TOF 5600 plus) and SWATH analysis was performed as described previously. Briefly, each sample was subjected to 1 IDA and 3 SWATH runs as technical replicates. IDA runs from MDA-468PC and MDA-468 PDTP-P were pooled together to generate an ion library and for analysis in Protein pilot software v4.5 (Sciex U.S.A) to obtain protein identities. No special factor was chosen. Digestion: Trypsin and cysteine alkylation: IAA, were the parameters set. The search was carried out against the Uniprot database containing human proteins as well as β gal from E.Coli. The proteins identified with 1% FDR were considered and the result file (.group) was generated. This result (.group) file served as the ion library. Peak View 2.2 software was used for SWATH run data extraction using the SWATH acquisition microapp (Sciex, USA). Proteins with 1% FDR identified from the spectral library were imported into peak View 2.2 software. The processing setting used was 6 peptides per protein, and 6 transitions per peptide. The peptide confidence threshold was set at 95% and the false discovery rate was chosen to be 5%. The retention time was calibrated. A common protein was selected with XIC (Extracted ion chromatogram) overlapping in all six SWATH injections. The processed data was saved as SWATH session file and the Ion library was saved as an updated RT.

The file was then exported into Marker view software, and the protein-based file was opened for further analysis. After normalization with cytosolic actin, PCA was applied where the samples were grouped as T (MDA MB 468 PDTP-P) and N (MDA-MB-468 PC). A t-test was conducted out to compare T to N to calculate the fold change and identify the up-regulated and down-regulated proteins.

### Immunoblotting

Whole cell lysate of different cell lines for total protein was prepared in lysis buffer consisting of 0.2% Triton X-100, 50 mM Hepes (pH 7.5), 100 mM NaCl, 1 mM MgCl2, 50 mM NaF, 0.5 mM NaVO3, 20 mM β-glycerophosphate, 1 mM phenylmethylsulfonyl fluoride, leupeptin (10 µg/ml), and aprotinin (10 µg/ml). The total protein concentration of the cell lysate was estimated by Bradford assay (Bio-Rad, Hercules, CA). Fifty micrograms of the total protein were used for SDS–polyacrylamide gel electrophoresis and transferred on nitrocellulose membrane, which proceeded as previously described method ((Verma et al. 2017)). The membrane was probed with Primary antibodies for STAT3, pSTAT3, AKT, pAKT, Src, Lyn, Fyn, EGFR, pEGFR, NDRG1, and GAPDH/beta-tubulin as a loading control.

### Cell viability assays MTT assay

To determine the cytotoxicity dose-response, experiments were performed in MDA-468 (12000 cells), HCC70 (9000 cells), and HS578T (3000 cells) using Gefitinib, Neratinib, and Paclitaxel. Cells were seeded in 96 well plates and were treated with drugs for 72 hrs. Cell viability was determined using MTT which was converted into blue formazan and solubilized. O.D was determined at 570 nm (reference wavelength 630 nm) using a spectrophotometer. Dose-response curves were generated using Graph-Pad Prism 5.0 software.

### Colony formation assay

Cells were seeded up to 50% confluency and treated with different concentrations of Gefitinib, Neratinib, and Paclitaxel for 72h to 96 hours. Media was replenished after every three days. The colony was fixed in 4% PFA and stained with crystal violet.

### Generation of 3D spheroids

3D spheroids were developed in U-shaped 96-well plate coated with 60µl of 1% agar in PBS in each well. 4000 cells (in 100 µl of medium) were seeded in each well and centrifuged at 1000g for 5 min. Plates were kept for 3 days in 5% CO2 humidified incubator. After three days, drug treatment was given for 15 days at an interval of 3 days. Phase contrast micrographs were acquired using 4X and 10X magnifications using an inverted microscope (OPTIKA, Italy).

### Immunofluorescence staining

Cells grown on coverslips were fixed in chilled methanol (10min at -20°C) and washed 3 times with PBS, followed by permeabilization with 0.3% triton in 0.1% NP40. Cells were incubated with primary antibody at 4°C overnight, followed by PBS and 0.1% NP40 alternate washes 3 times. Alexa 488 and 560 Secondary antibody incubation was performed for 1 hr at room temperature. Nuclear staining was done with DAPI.

### Bioinformatic analysis

#### Datasets

Gene and protein expression and correlation analysis of breast cancer patients were done based on The Cancer Genome Atlas (TCGA) and protein atlas (TCPA) breast cancer dataset. For differential gene expression in TNBC subtypes and DEPMAP analysis UALCAN platform (https://ualcan.path.uab.edu/analysis.html) was used. For Survival analysis, TCGA gene expression was used on Kaplan-Meier Plotter (https://kmplot.com/analysis) and Recurrence Free Survival (RFS) was analyzed using median cut. For protein expression TCPA portal (https://tcpaportal.org/tcpa/survival_analysis.html) was used to derive survival, and Kaplan-Meier plots were derived with BRCA data set containing 901 patient samples. ROC plots in chemotherapy-treated patients were generated using the ROC plotter portal (http://www.rocplot.org/site/index) (Fekete and Győrffy 2019).

### Statistical analysis

Data are presented as mean ±SD. To compare between experimental groups, we used Student’s t-test for experiments. The reported p-values associated with dataset analysis (TCGA or TCPA) are generated using online portals. p-value<0.05 is considered significant.

## Acknowledgments

This research work was supported by grants from Basic and Translational Research in Cancer grant (No.1/3(7)/2020/TMC/R&D-II/8823 Dt.30.07.2021), and Animal Imaging at ACTREC, grant No. 1/3(6)/2020/TMC/R&D -II/ 3805 from Department of Atomic Energy (DAE), Government of India.

## Authors’ contributions

NC: designed the study, performed experiments, analyzed the data, and written parts of the manuscript; BSC: designed and performed the experiments, analyzed data, and written parts of the manuscript; AS, SM, and KV: designed and performed experiments, analyzed data; SES prepared the samples for mass spectrometry analysis and written parts of manuscript; SK performed the analysis of mass spectrometry data; JB and JD: performed parts of WB analysis, JD performed the densitometric analysis of WB. NV: designed the conception, developed experimental methods and models, acquired funding, performed and analyzed experimental and in-silico data, supervised and compiled the study, and prepared the manuscript. All authors read and approved the final manuscript.

