## Supplementary material for "EGFR-to-Src family tyrosine kinase switching in proliferating-DTP TNBC cells creates a hyperphosphorylation-dependent vulnerability to EGFR TKI"

***Running title:*** *EGFR-Src kinase switching creates vulnerability to EGFR-TKi in TNBC DTPs*

**Supplementary Table 1.** Differentially expressed cellular proteins between MDA-MB-468 PC and PDTP-P as analyzed by Mass spectrometry analysis.

| S.No. | Protein | Protein_abv. | Fold_Change (FC) | log2fc | padj |
|---|---|---|---|---|---|
| **1** | **Epidermal growth factor receptor** | **EGFR** | **3.52412968829552E-2** | **-4.82659** | **2.16E-05** |
| **2** | Protein NipSnap homolog 2 | NIPSNAP2 | 6.3380510263221695E-2 | -3.97982 | 0.00027 |
| **3** | D-3-phosphoglycerate dehydrogenase | PHGDH | 8.3938999433552999E-2 | -3.57451 | 0.015116 |
| **4** | 40S ribosomal protein S18 | RPS18 | 8.6851150959902104E-2 | -3.52531 | 0.004865 |
| **5** | Keratin, type II cytoskeletal 8 | KRT8 | 9.2480378932206203E-2 | -3.43471 | 0.000511 |
| **6** | 40S ribosomal protein S15a | RPS15A | 9.5235264545156295E-2 | -3.39236 | 0.016238 |
| **7** | Proliferation-associated protein 2G4 | PA2G4 | 9.7442002172023298E-2 | -3.35931 | 0.000313 |
| **8** | 60S ribosomal protein L27a | RPL27A | 0.108458764580646 | -3.20478 | 0.006654 |
| **9** | Keratin, type I cytoskeletal 19 | KRT19 | 0.113852863833228 | -3.13476 | 0.008901 |
| **10** | 60S ribosomal protein L4 | RPL4 | 0.138041977980004 | -2.85682 | 0.005261 |
| **11** | 60S ribosomal protein L7 | RPL7 | 0.151450509387549 | -2.72308 | 0.001461 |
| **12** | Malate dehydrogenase, cytoplasmic | MDH1 | 0.15922759885092799 | -2.65084 | 0.029407 |
| **13** | Keratin, type II cytoskeletal 5 | KRT5 | 0.16439139458433699 | -2.60479 | 0.000654 |
| **14** | Keratin, type I cytoskeletal 14 | KRT14 | 0.17467792362905199 | -2.51723 | 0.012478 |
| **15** | Keratin, type I cytoskeletal 16 | KRT16 | 0.19241348374249501 | -2.37772 | 0.010415 |
| **16** | 40S ribosomal protein S14 | RPS14 | 0.19551679330100899 | -2.35464 | 0.029247 |
| **17** | Cytochrome c | CYCS | 0.198803358388639 | -2.33059 | 0.012478 |
| **18** | Heterogeneous nuclear ribonucleoprotein Q | SYNCRIP | 0.209348361334457 | -2.25602 | 0.024988 |
| **19** | 40S ribosomal protein S2 | RPS2 | 0.21953854623000299 | -2.18745 | 0.013128 |
| **20** | Platelet-activating factor acetylhydrolase IB subunit gamma | PAFAH1B3 | 0.23534305402883801 | -2.08716 | 0.020659 |
| **21** | Keratin, type I cytoskeletal 17 | KRT17 | 0.25197413376908101 | -1.98865 | 0.005299 |
| **22** | 60S ribosomal protein L6 | RPL6 | 0.25286964853163701 | -1.98353 | 0.029247 |
| **23** | Histone H1.5 | H1-5 | 0.26591043730578301 | -1.91099 | 0.00513 |
| **24** | 60S ribosomal protein L10-like | RPL10L | 0.26921167527970302 | -1.89319 | 0.022692 |
| **25** | Non-POU domain-containing octamer-binding protein | NONO | 0.28693714203256698 | -1.80119 | 0.013581 |
| **26** | Histone H4 | H4C1 | 0.28710566743875099 | -1.80035 | 0.000313 |
| **27** | 40S ribosomal protein S4, X isoform | RPS4X | 0.29046369044824999 | -1.78357 | 0.001423 |
| **28** | Serine hydroxymethyltransferase, mitochondrial | SHMT2 | 0.30445623900276603 | -1.71569 | 0.037668 |
| **29** | Heat shock protein HSP 90-alpha | HSP90AA1 | 0.31639649502600398 | -1.66019 | 0.02346 |
| **30** | Histone H2A.J | H2AFJ | 0.32677032744643603 | -1.61365 | 0.002506 |
| **31** | GTP-binding nuclear protein Ran | RAN | 0.33326596957029098 | -1.58525 | 0.001239 |
| **32** | Elongation factor 1-gamma | EEF1G | 0.33782658440915397 | -1.56565 | 0.012478 |
| **33** | Keratin, type II cytoskeletal 6C | KRT6C | 0.33933056038919301 | -1.55924 | 0.013581 |
| **34** | Isoform 2 of Histone H2B type 2-F | HIST2H2BF | 0.35071635750372199 | -1.51162 | 0.00027 |
| **35** | 60 kDa heat shock protein, mitochondrial | HSPD1 | 0.35560604371916099 | -1.49165 | 0.002185 |
| **36** | Proliferating cell nuclear antigen | PCNA | 0.37625846388086198 | -1.4102 | 0.013128 |
| **37** | Phosphoglycerate kinase 1 | PGK1 | 0.38636598040678699 | -1.37196 | 0.009081 |
| **38** | ATP synthase subunit alpha, mitochondrial | ATP5F1A | 0.39693128987197401 | -1.33304 | 0.012232 |
| **39** | ADP/ATP translocase 2 | SLC25A5 | 0.39871555505983503 | -1.32657 | 0.020534 |
| **40** | Fructose-bisphosphate aldolase A | ALDOA | 0.410866131826777 | -1.28326 | 0.016238 |
| **41** | 60S ribosomal protein L12 | RPL12 | 0.48606808542914998 | -1.04077 | 0.000654 |
| **42** | Elongation factor 1-alpha 1 | EEF1A1 | 0.48877493673023897 | -1.03276 | 0.00027 |
| **43** | L-lactate dehydrogenase A chain | LDHA | 0.49017431658405197 | -1.02863 | 0.028049 |
| **44** | Tubulin beta chain | TUBB | 0.49610708339903198 | -1.01128 | 0.027471 |
| **45** | Prelamin-A/C | LMNA | 0.50489136546989499 | -0.98596 | 0.025276 |
| **46** | Glyceraldehyde-3-phosphate dehydrogenase | GAPDH | 0.50670297398284503 | -0.98079 | 0.015116 |
| **47** | Triosephosphate isomerase | TPI1 | 1.5579341820314501 | 0.639634 | 0.007709 |
| **48** | Peroxiredoxin-5, mitochondrial | PRDX5 | 1.985495404064 | 0.989499 | 0.030144 |
| **49** | Nucleophosmin | NPM1 | 2.21836198677833 | 1.149495 | 0.038156 |
| **50** | Myosin light polypeptide 6 | MYL6 | 2.72845765778988 | 1.448086 | 0.00513 |
| **51** | Cathepsin D | CTSD | 2.8151734049180202 | 1.493224 | 0.024802 |
| **52** | Beta-galactosidase | GLB1 | 2.95111751784275 | 1.561261 | 0.025375 |
| **53** | Alpha-2-HS-glycoprotein | AHSG | 3.0912093838344599 | 1.628171 | 0.006654 |
| **54** | SH3 domain-binding glutamic acid-rich-like protein 3 | SH3BGRL3 | 4.2990086818932802 | 2.104004 | 0.00613 |
| **55** | Lactotransferrin | LTF | 5.3390460132630198 | 2.416582 | 8.08E-05 |
| **56** | Vitamin D-binding protein | GC | 10.1873566862981 | 3.348708 | 0.001122 |
| **57** | Serum albumin | ALB | 11.1437016335029 | 3.478157 | 8.08E-05 |
| **58** | Afamin | AFM | 11.815142958116899 | 3.562565 | 0.00513 |
| **59** | Galectin-1 | LGALS1 | 17.8344009472567 | 4.156591 | 0.000145 |

Figure 1S

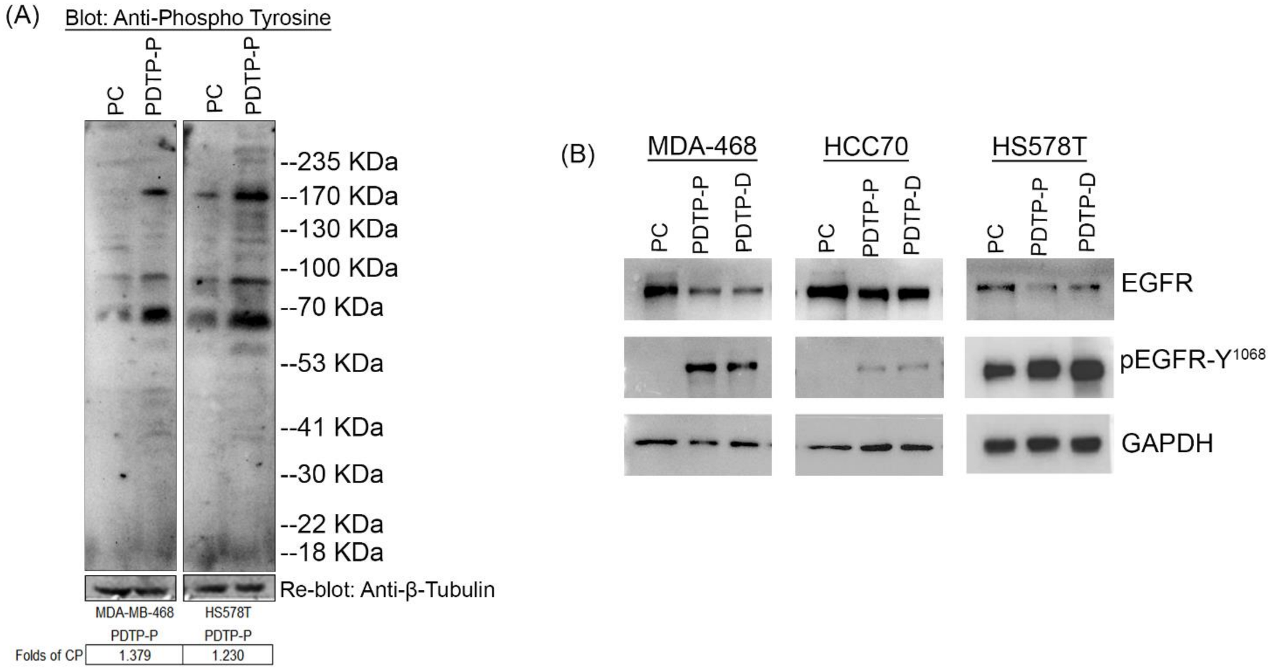


**Figure 1S. (A)** Phospho-tyrosine signal was detected in the whole cell lysates of MDA-468 and HS578T chemotherapy naïve parental cells (PC) and paclitaxel-derived proliferating drug-tolerant (PDTP-P) cells by Western blot (WB) analysis using an anti-phospho-tyrosine antibody. The membrane was re-blotted with anti-β-tubulin antibody to indicate the loading control. Molecular weight marker sizes are indicated. The mean intensity of the phospho-tyrosine signal in the whole lanes was quantified densitometric analysis and the folds change to PC was calculated for PDTP cells and indicated below the WB. **(B)** Levels of EGFR and p-EGFR were determined in different subtypes of TNBC cell lines and their Paclitaxel- or Doxorubicin-derived PDTPs as assessed by WB analysis. GAPDH served as their loading control. The expression levels of the indicated proteins.

Figure S2

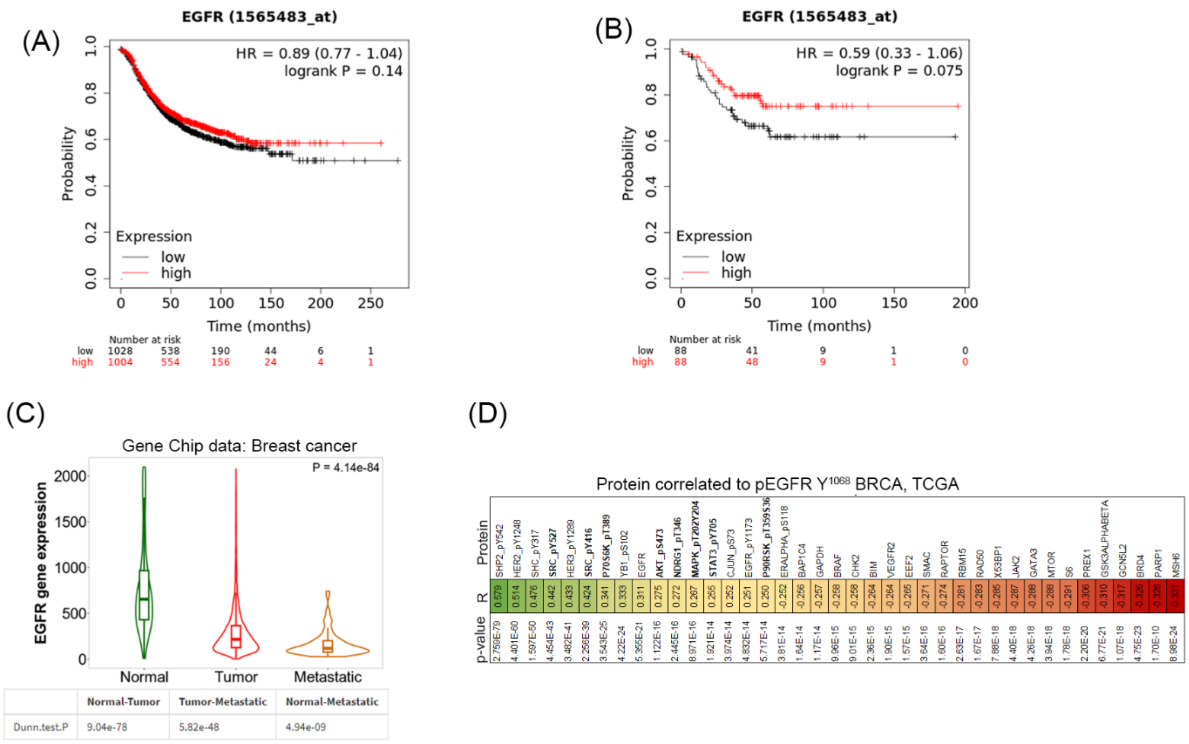


**Figure S2: (A)** Kaplan-Meier curve from untreated breast cancer patient TCGA cohort showing RFS in patients with mRNA expression of EGFR tumor tissue stratified into a high or low expression using the median expression value as the cut-off point. The corresponding P-value for Log-rank test in all untreated BC patients was shown. No change between low and high EGFR levels. Similarly, **(B)** Kaplan-Meier curve showing RFS in patients with mRNA expressions of EGFR in untreated TNBC that were stratified into high or low expression using the median expression value as the cut-off point. The corresponding P-value for Log-rank test in TNBC irrespective of systemic treatments. **(C)** The violin plot showing EGFR gene expression in normal human breast tissues, in breast cancer tissue, and in metastatic breast cancer tissues. **(D)** Heatmap showing Spearman's rank correlation coefficient (R) for proteins significantly correlated with expression of pEGFR $Y^{1068}$ using proteomics data available at the TCPA (The Cancer Protein Atlas) from the TCGA-BRCA cohort (901 patients).

Figure S3

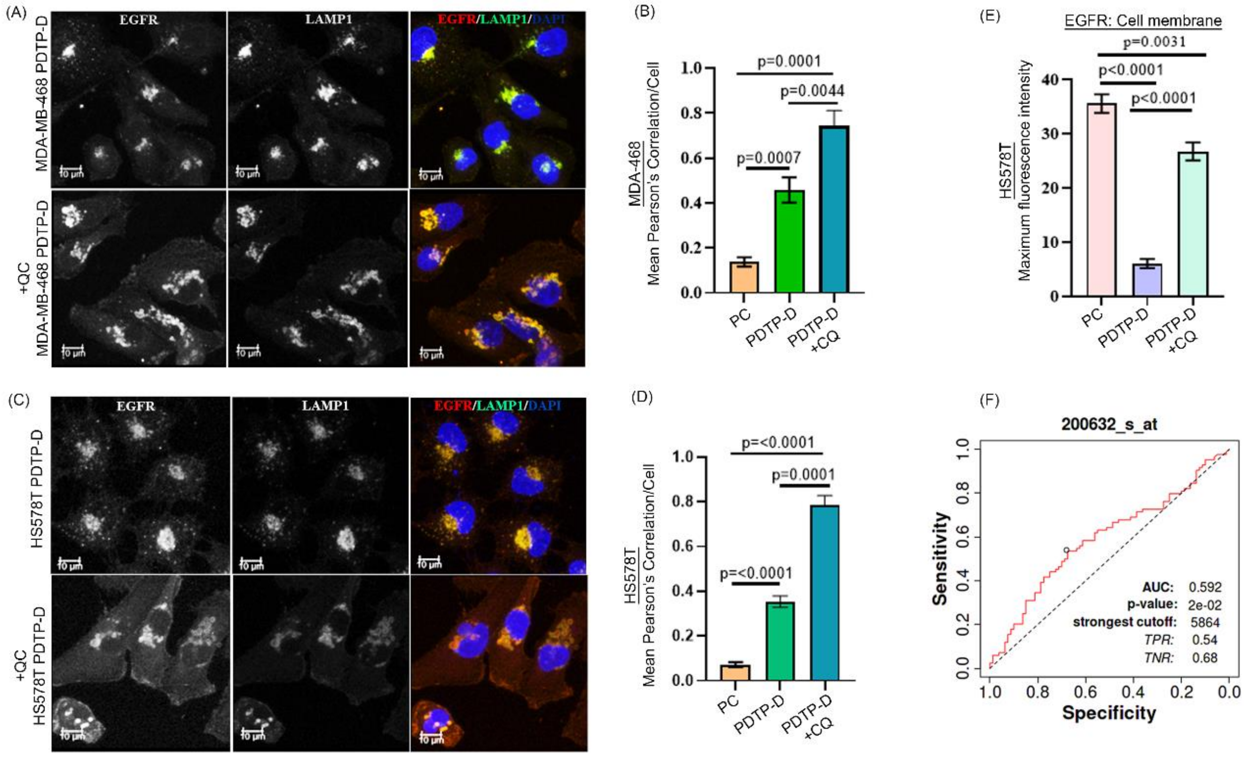


**Figure S3: (A)** and **(C)** Representative confocal microscope images of the parental TNBC cell lines (PC), MDA-468 and HS578T (PC) and Doxorubicin derived proliferating drug-tolerant persister (PDTP-D) cells stained with EGFR, LAMP1, and DAPI with and without 25 µM of CQ treatment for 12 hrs as described in Materials and Methods. Scale bar, 10 µm. **(B) and (D)** Bar graphs showing the quantified and analyzed mean colocalization coefficient for EGFR (RED channel) and LAMP1 (Green channel) in TNBC PC and PDTPs from immunofluorescence staining analysis shown in A and C. **(E)** Bar plot showing cell surface expression of EGFR in HS578T PC and PDTP-P cells and its rescue upon treatment with CQ, quantified from the immunofluorescence attaining analysis shown in C. **(F)** ROC curve comparing sensitivity and specificity of NDRG1 gene for classifying responders vs. non-responders to the chemo treatment in all TNBC chemo treated patient samples showing 5-year recurrence-free survival.

Figure S4

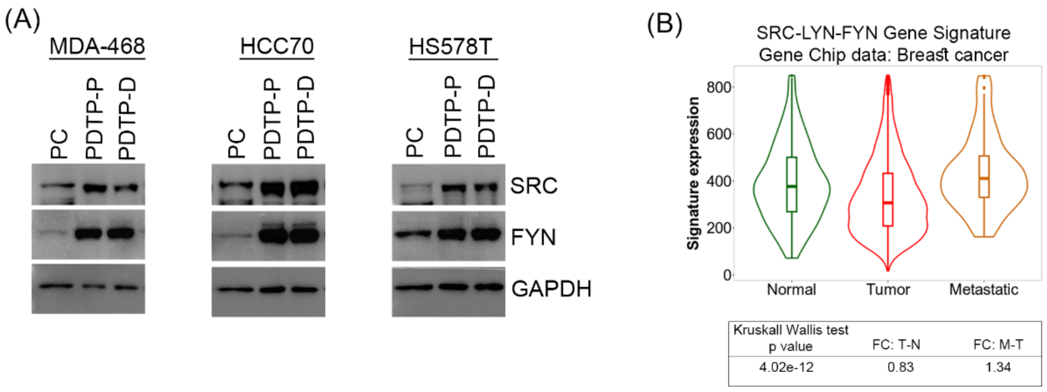

**Figure S4:** Levels of SRC and FYN were determined in Different subtypes of TNBC PC and their Paclitaxel- and Doxorubicin-derived PDTPs. GAPDH served as their loading control. The expression levels of the indicated proteins were assessed by WB, and band intensities were quantified by ImageJ software and presented as fold of control in the bar graph. **(B)** Violin plot showing expression of a triple gene signature SRC-LYN-FYN in normal breast tissue, in primary breast tumor and in metastatic tumor samples in human tissues cohort in TCGA. Table below the graph indicating the p-value and fold change (FC) in gene expression between tumor vs normal tissues (T-N) and metastatic vs primary tumor tissues (M-T). Krushkall Wallis test was applied to calculate the variance between the groups and p-value for significance.

Figure S5

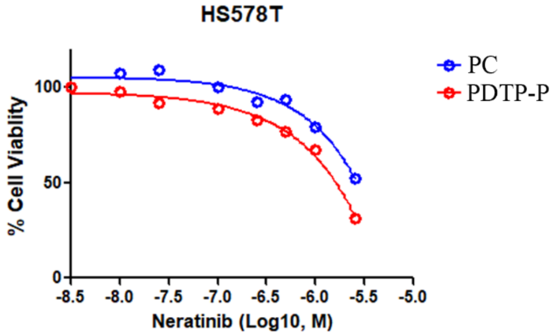


**Figure S5:** Dose-response curves of varying concentrations Gefitinib in the PC HS578T cell line and its Paclitaxal PDTP treated for 72 hours. Dose-response curves are presented as means of three repeats.

Figure S6

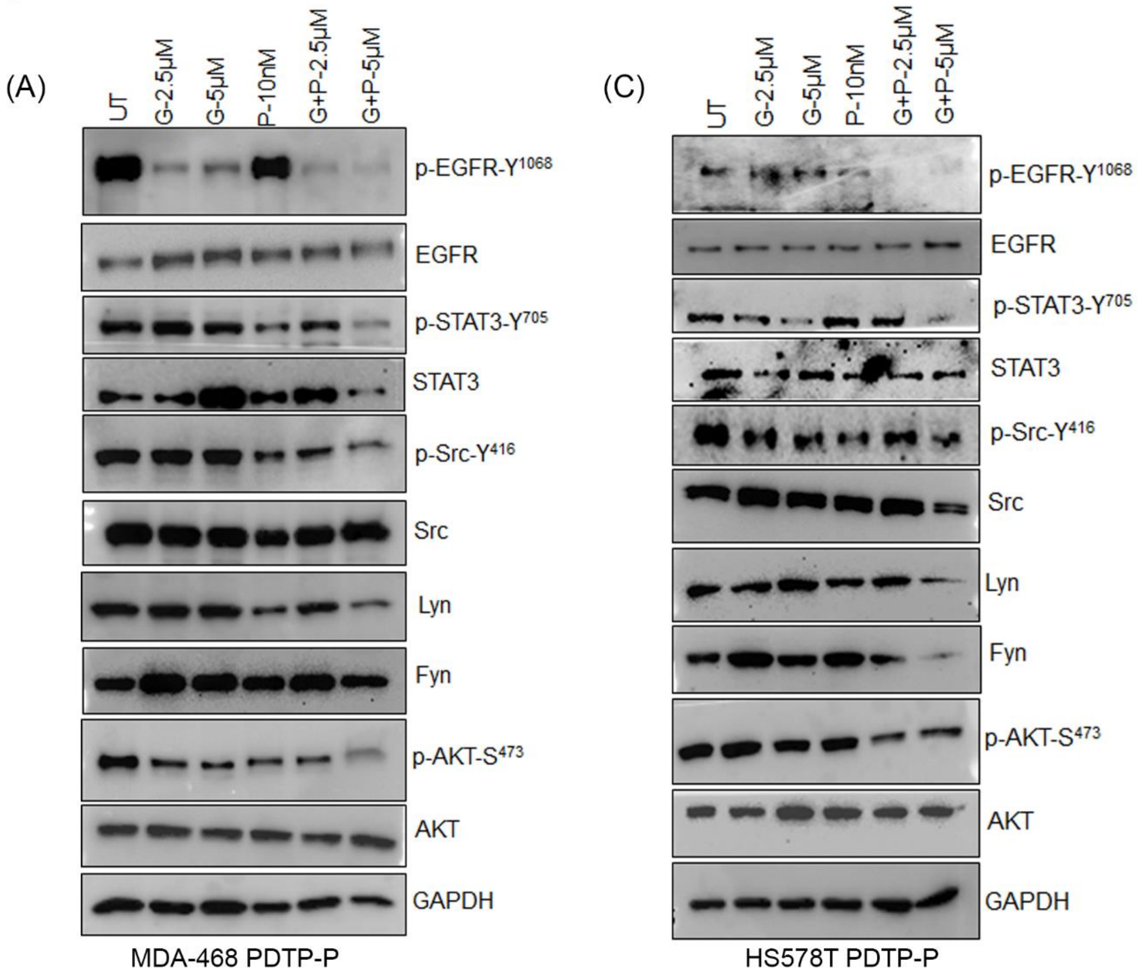


(B)

| | pEGFR | pSRC | pSTAT3 | pAKT | SRC | FYN | LYN |
|---|---|---|---|---|---|---|---|
| Untreated | 1.00 | 1.00 | 1.00 | 1.00 | 1.00 | 1.00 | 1.00 |
| Gef. 2.5µM | 0.41 | 0.95 | 0.99 | 0.70 | 1.04 | 1.01 | 1.01 |
| Gef. 5µM | 0.24 | 0.96 | 0.99 | 0.57 | 1.07 | 0.92 | 1.00 |
| Paclitaxel 10nM | 0.56 | 0.84 | 0.78 | 0.75 | 0.86 | 0.67 | 0.91 |
| Gef. 2.5µM+Paclitaxel | 0.49 | 0.57 | 0.67 | 0.69 | 0.89 | 0.58 | 0.71 |
| Gef. 5µM+Paclitaxel | 0.21 | 0.38 | 0.57 | 0.48 | 0.83 | 0.58 | 0.54 |

MDA-MB-468 PDTP-P

(D)

| | pEGFR | pSRC | pSTAT3 | pAKT | SRC | FYN | LYN |
|---|---|---|---|---|---|---|---|
| Untreated | 1.00 | 1.00 | 1.00 | 1.00 | 1.00 | 1.00 | 1.00 |
| Gef. 2.5µM | 0.91 | 0.89 | 0.69 | 1.05 | 0.92 | 1.01 | 1.01 |
| Gef. 5µM | 0.91 | 0.94 | 0.52 | 1.02 | 0.99 | 0.92 | 1.00 |
| Paclitaxel 10nM | 0.72 | 0.86 | 0.77 | 1.05 | 0.96 | 0.67 | 0.91 |
| Gef. 2.5µM+Paclitaxel | 0.60 | 0.77 | 0.62 | 0.64 | 0.96 | 0.58 | 0.71 |
| Gef. 5µM+Paclitaxel | 0.59 | 0.58 | 0.40 | 0.59 | 0.52 | 0.58 | 0.54 |

HS578T PDTP-P

**Figure 6.** Western blot analysis to assess the proteomic changes in PDTP-P of MDA-468 **(A)** and HS578T **(B)** in response to Gefitinib, paclitaxel single or combined treatments after 24 h as determined by probing the total cellular lysate with the indicated antibody. **(B)** and **(D)** ImageJ software was used for quantification of Western blot band intensities of PC and PDTP-P in A and C and presented as heat map of changes folds of untreated PC in each type.
